# Pyruvate oxidase as a key determinant of pneumococcal viability during transcytosis across the blood-brain barrier endothelium

**DOI:** 10.1101/2020.12.31.424967

**Authors:** Anjali Anil, Akhila Parthasarathy, Shilpa Madhavan, Kwang Sik Kim, Anirban Banerjee

## Abstract

*Streptococcus pneumoniae* (SPN / pneumococcus), invades myriad of host tissues following efficient breaching of cellular barriers. However, strategies adopted by pneumococcus for evasion of host intracellular defences governing successful transcytosis across host cellular barriers remain elusive. In this study, using brain endothelium as a model host barrier, we observed that pneumococcus containing endocytic vacuoles (PCVs) formed following SPN internalization into brain microvascular endothelial cells (BMECs), undergo early maturation and acidification, with a major subset acquiring lysosome-like characteristics. Exploration of measures that would preserve pneumococcal viability in the lethal acidic pH of these lysosome-like vacuoles revealed a critical role of the two-component system response regulator, CiaR, which has been previously implicated in induction of acid tolerance response. Pyruvate oxidase (SpxB), a key sugar metabolizing enzyme that catalyses oxidative decarboxylation of pyruvate to acetyl phosphate, was found to contribute to acid stress tolerance, presumably via acetyl phosphate-mediated phosphorylation and activation of CiaR, independent of its cognate kinase CiaH. Hydrogen peroxide, the by-product of SpxB catalysed reaction, was also found to improve pneumococcal intracellular survival, by oxidative inactivation of lysosomal cysteine cathepsins, thus compromising the degradative capacity of the host lysosomes. Expectedly, a Δ*spxB* mutant was found to be significantly attenuated in its ability to survive inside the BMEC endocytic vacuoles, reflecting in its reduced transcytosis ability. Collectively, our studies establish SpxB as an important virulence determinant facilitating pneumococcal survival inside host cells, ensuring successful trafficking across host cellular barriers.

**AUTHOR SUMMARY:** Eukaryotic cells which constitute host barriers have innate immune defences to restrict microbial passage into sterile compartments. This necessitates need for pathogens to devise strategies to evade these, for successful establishment of disease. In this study, by focussing on the blood-brain barrier endothelium, we investigate the mechanisms which enable the opportunistic pathogen *Streptococcus pneumoniae* to traverse host barriers. Pyruvate oxidase, a pneumococcal sugar metabolizing enzyme was found to play a critical role in this key event, owing to production of acetyl phosphate and hydrogen peroxide via its enzymatic activity. On one hand, acetyl phosphate, by contributing to activation of acid tolerance stress response, enabled pneumococci to maintain viability in the lethal acidic pH of the lysosome-like vacuoles inside brain endothelium. On the other, hydrogen peroxide, was found to oxidise and inactivate a subset of degradative lysosomal enzymes. This two-pronged approach, aided by pyruvate oxidase, enabled pneumococci to evade intracellular degradation for successful transcytosis across the endothelium. Thus, pyruvate oxidase is a key determinant of pneumococcal virulence and hence can potentially serve as a viable candidate for therapeutic interventions for better management of invasive pneumococcal diseases.

## INTRODUCTION

A constant threat of attack by pathogenic microorganisms has driven evolution to refurbish several of the eukaryotic homeostasis pathways for defence against invading microbes and maintain the sanctity of the intracellular space. These include the endocytic pathway, autophagy and ubiquitin-proteasome machinery and these are appropriate choices owing to their role in diverse bio-molecular degradations. As a consequence, for successful establishment of disease, pathogens must employ strategies to prevent their degradation in the intracellular milieu. This is especially crucial for intracellular pathogens such as *Mycobacterium tuberculosis, Salmonella typhimurium, Legionella pneumophila* etc. Interestingly, in recent times, bacterial pathogens that were previously categorized as extracellular, have been found to replicate in certain host cell types, examples of which include *Streptococcus pyogenes* and *Staphylococcus aureus* [1–3]. An intracellular niche, especially within non-professional phagocytes has been proposed to act as reservoirs for persistent or latent infections and provide sanctuary from the extracellular foes such as complement system and antibiotics. Thus, outcome of the tug-of-war between invading pathogen and host intracellular defences is an important milestone that governs the trajectory of infection.

*Streptococcus pneumoniae* (SPN / pneumococcus), the opportunistic pathogen, causes life threatening diseases such as pneumonia, bacteremia, endocarditis and meningitis. Although typically considered an extracellular pathogen, it does have a brief intracellular stint during trafficking across host barriers, mainly the blood-brain barrier (BBB). Recent studies have also identified that SPN is capable of prolonged residence and replication inside splenic macrophages, serving as transient reservoirs for dissemination into the blood causing septicaemia [4]. Additionally, SPN has been shown to replicate within cardiomyocytes [5], Pneumococcal factors such as the polysaccharide capsule, choline binding proteins (PspA, PspC), exoglycosidases (NanA, BgaA, StrH) and IgA1 protease that contribute towards its immune evasion has been extensively investigated [6–9]. However, the strategies that enable SPN to evade diverse host intracellular defences remain elusive.

In this study, we tracked the fate of Pneumococcus Containing endocytic Vacuoles (PCVs) formed following SPN internalization into human brain microvascular endothelial cells (hBMECs), the primary constituent of BBB. Neurotropism of SPN predominantly results from its intimate interaction with the brain endothelium which it breaches primarily by adopting the transcytosis route [10], a mode of passage necessitating a brief but stable intracellular life, for efficient BBB traversal. Pneumococcal invasion of the BBB endothelium is facilitated via interaction of its surface molecules with endothelial cell receptors (reviewed in [11]), in turn inducing its internalization via multiple endocytic routes including clathrin and caveolae-dependent and dynamin-independent pathways [12]. However, irrespective of the internalization mode, SPN has been shown to interact with and get degraded via xenophagy and ubiquitin-proteasome machinery [13]. Although a large fraction of internalized pneumococci gets degraded during BBB transit, for establishment of a successful infection it is a prerequisite that a subset of SPN successfully transcytose through the BMECs and enter the brain. Although few pneumococcal features including the pili, pneumolysin expression, phase and chain size has been implicated in governing the fate of BMEC-invaded SPN [13–15], detailed mechanisms remain unexplored.

In this study we observed that following invasion, a major fraction of PCVs mature and get acidified very early on, with a major subset also showing association with the lysosomal marker cathepsin B. In exploring the factors that enable SPN to remain viable within the acidified lysosome-like compartments, we identified that the activity of the two-component system response regulator CiaR is critical and foster pneumococcal survival by activating the acid tolerance stress response. Interestingly, the sugar metabolizing enzyme pyruvate oxidase (SpxB) which converts pyruvate to acetyl phosphate was found to play an important role in inducing the acid tolerance response, via acetyl phosphate-mediated activation of CiaR, independent of its cognate kinase CiaH. SpxB mutants were found to be significantly attenuated in survival under lethal acidic stress *in vitro* as well as within the hBMEC vacuoles. Additionally, the by-product of SpxB catalysed reaction, hydrogen peroxide, was found to prolong pneumococcal survival inside hBMECs, by interfering with the proteolytic activity of lysosomal cysteine cathepsins. Thus, we implicate acid stress tolerance as a survival strategy and pyruvate oxidase as a key factor governing the viability of SPN during trafficking across host barriers such as the BBB.

## RESULTS

### Pneumococci persists within acidified vacuoles during transit through brain endothelium

Although microbes exploit eukaryotic endocytosis as a portal for entry into host cells, the pathway act as the primary line of defence against invading microbes, owing to its role in intracellular degradation. Hence, we first sought to understand the trajectory of pneumococcus containing endocytic vacuoles following entry into the hBMECs. Typically, the endocytic vacuole formed following uptake of extracellular cargo undergoes a maturation process involving changes in the lipid and protein composition of the vacuolar membrane and progressive acidification of the vacuolar lumen [16]. The vacuole formed initially, called the early endosome (marked by phosphatidylinositol 3-phosphate (PI3P) and the GTPase - Rab5) [17] subsequently matures to form late endosomes (marked by phosphatidylinositol 3, 5-bisphosphate (PI3,5P_2_) and Rab7), finally fusing with lysosomes resulting in degradation of the cargo, brought about by the action of plethora of hydrolases that function optimally at acidic pH (pH ~ 4.5) [16]. To assess how SPN has modified the vacuolar identity for improved intracellular survival, association of PCVs with these different endosomal markers were investigated by immunofluorescence microscopy.

Immediately following internalization, at 0 hour post infection (h.p.i), 42.07 ± 2.544% of wild type (WT) SPN vacuoles were found to be positive for P40PX-EGFP (Phox homology domain which binds to PI3P) [18, 19], confirming its presence within early endosomal vacuoles (Fig 1A, B). Next, we checked for acidification of PCVs, using a fluorescent acidotropic dye, Lysotracker. Time-course analysis of SPN co-localization with Lysotracker revealed that vast majority of PCVs get acidified as early as 2 h.p.i. (52.4 ± 5.177%), whose proportion remain similar till later time points (45.08 ± 2.126% at 6 h.p.i and 42.3 ± 5.089% at 10 h.p.i) (Fig 1C, D). A fraction of PCVs also showed characteristics of fusion with lysosomes, marked by their co-localization with cathepsin B (12.8 ± 1.555% at 10 h.p.i), one of the most abundant lysosomal proteases, belonging to the family of cysteine cathepsins [20] (Fig 1E, F). Overall, these suggest that majority of PCVs undergo classical endocytic maturation leading to acidification and subsequent fusion with lysosomes.

**Fig 1.**
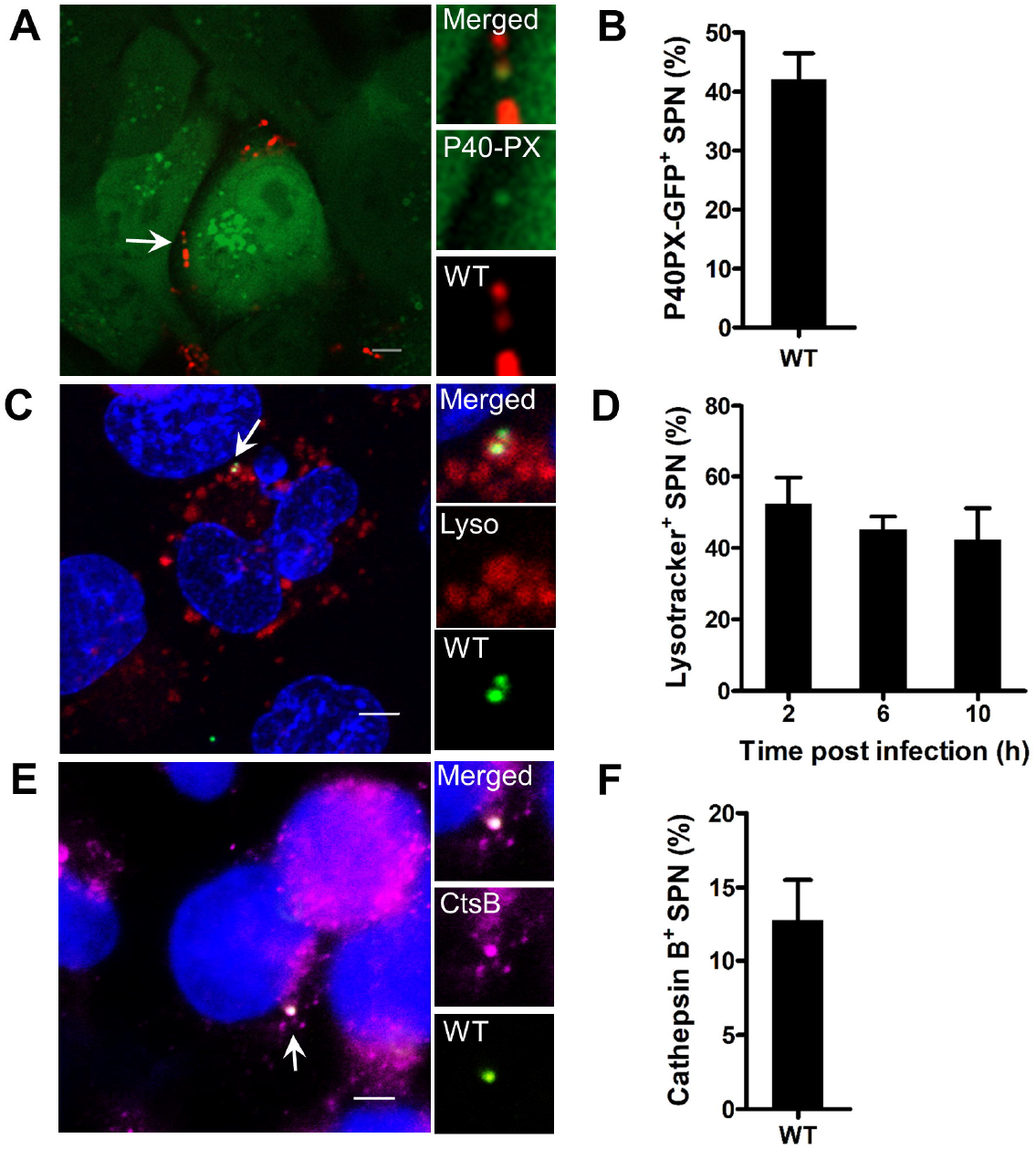
Pneumococci persists within acidified vacuoles of the brain endothelium. Confocal micrographs showing association of (**A**) SPN-tagRFP (red) with P40PX-EGFP (green) at 0 h.p.i. (**C**) SPN-GFP (green) with Lysotracker (Red) at 6 h.p.i. (**E**) SPN (green) with cathepsin B (pink) at 10 h.p.i. DAPI (blue) has been used to stain hBMEC nucleus in (C) and (E). Arrows designate the bacteria shown in insets. Event was localized at z-stack number: 1-2 out of 17 (for A), 2-3 out of 6 (for C) and 9 out of 17 (for E). (**B**) Quantification of co-localization of WT SPN with the early endosomal marker P40PX-EGFP, at 0 h.p.i in hBMECs (**D**) Time course analysis of association of WT SPN with Lysotracker (a marker for acidified vacuoles) in hBMECs. (**F**) Quantification of co-localization of WT SPN with the lysosomal marker cathepsin B, at 10 h.p.i in hBMECs. Data information: Data is presented as mean ± SD of triplicate wells. n ≥ 100 SPN per coverslip (B, D, F).

### Pyruvate oxidase contributes to pneumococcal acid tolerance response, via regulation of CiaR

Significant association of PCVs with Lysotracker at all time points suggested that SPN must own strategies to withstand the acidic stress in order to remain viable for successful transcytosis across the brain endothelium. Previous studies have revealed that SPN possess an acid tolerance response (ATR), which enables it to survive exposure to lethal pH (4.4) if pre-exposed to a sub-lethal pH [21]. The gradual drop in pH of the vacuolar lumen, during its maturation represents a similar scenario, wherein SPN residence within the early and late endosomal vacuoles (whose pH is sub-lethal), can potentially enable it to tolerate the lethal pH of the lysosomes, thereby maintaining its viability. SPN possess an impressive assortment of two-component systems (TCSs) which equip it to sense and respond appropriately to environmental signals. Among these, the CiaRH TCS (competence induction and altered cefotaxime susceptibility, tcs05, CiaH-histidine kinase and CiaR-response regulator) [22], has been implicated in ATR induction, promoting pneumococcal survival in acidic conditions *in vitro* [23].

In order to check whether the CiaRH TCS plays a role in pneumococcal survival within the brain endothelium, we performed intracellular survival assay using WT and Δ*ciaR* mutant strains wherein the latter demonstrated attenuated survival from 4 h.p.i onwards (S1A Fig). A non-phosphorylatable CiaR mutant (R6:CiaR^D51A^), mimicked the phenotype of Δ*ciaR* mutant (S1A Fig), indicating that phosphorylation and subsequent activation of CiaR allowing it to regulate the expression of genes under the CiaR regulon, is necessary for SPN survival inside hBMECs, possibly by enabling it to mount the ATR. The contribution of CiaRH TCS and specifically, CiaR response regulator activity, for pneumococcal survival under lethal acidic stress was additionally confirmed by an *in vitro* ATR assay (S1B, C Fig).

Since CiaH is the cognate kinase for CiaR, we next compared the intracellular survival capability of Δ*ciaR* and Δ*ciaH* mutant strains in hBMECs. Although, Δ*ciaH* demonstrated reduced survival capability when compared to WT (S1D Fig), it was significantly higher than that of the Δ*ciaR* mutant (~ 2.5-fold at 8 h.p.i) (Fig 2A). This suggested the existence of another moiety that can phosphorylate and activate CiaR, in turn facilitating pneumococcal survival inside the host cells. Historically, acetyl phosphate (AcP) has been used as a phosphate donor for *in vitro* phosphorylation of purified response regulators, including CiaR [22]. Very recently, AcP was confirmed to independently phosphorylate CiaR in SPN [24]. Interestingly, AcP is formed as a result of sugar metabolism in SPN in a reaction catalysed by the enzyme pyruvate oxidase (SpxB). SpxB catalyses the oxidative decarboxylation of pyruvate to AcP, generating hydrogen peroxide as a by-product [25]. In order to investigate if SpxB contribute to ATR induction, we first compared the viability of Δ*ciaH* mutant with that of a Δ*ciaH*Δ*spxB* double mutant in an *in vitro* ATR assay. We observed a significant reduction in viability for the latter (~ 2-fold) suggesting that SpxB contributes to acid stress tolerance in SPN (Fig 2B). To confirm that the reduction in viability of the Δ*ciaH*Δ*spxB* double mutant is indeed due to loss of SpxB-mediated activation of CiaR, we constructed a reporter strain SPN:*P_htrA_*-GFP. This strain has GFP cloned under the 5’ untranslated region (5’UTR) of *htrA*, one of the genes strongly (positively) and solely regulated by CiaR [26]. Comparison of GFP expression in SPN:*P_htrA_*-GFP variants: WT, Δ*ciaH* and Δ*ciaH*Δ*spxB*, by flow cytometry, revealed a gradation in GFP expression, with the double mutant exhibiting the lowest; both in terms of GFP positive population (Fig 2C) and its mean fluorescence intensity (Fig 2D). Next, we subjected these strains to progressively reducing pH; from 7.1 to 5.9 (sub-lethal) and then to 4.4 (lethal) and observed that the GFP positive population was lowest in the Δ*ciaH*Δ *spxB* double mutant, even under acidic stress (S1E Fig).

**Fig 2.**
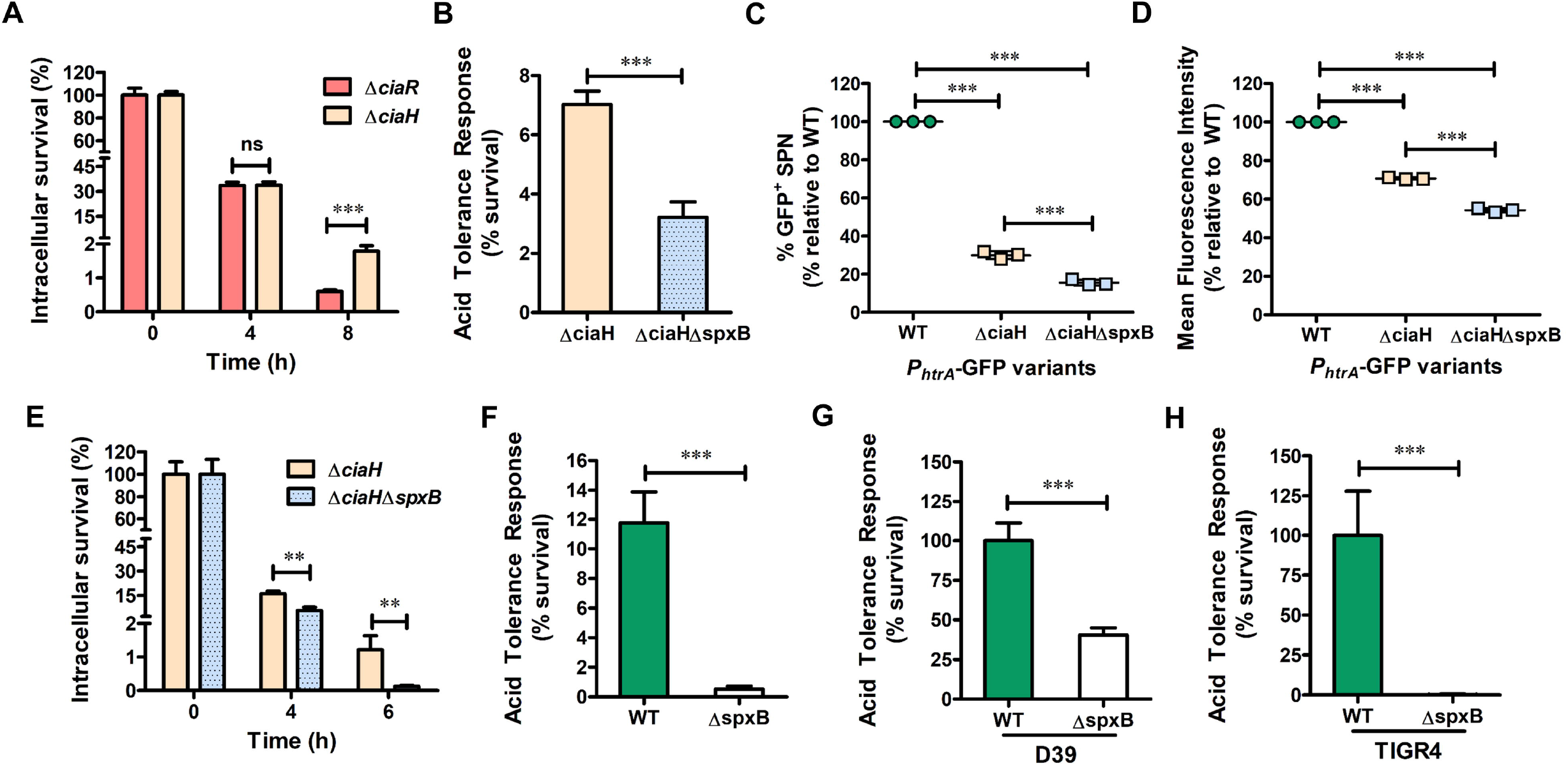
Pyruvate oxidase contributes to pneumococcal acid tolerance response, via regulation of CiaR. (**A**) Intracellular survival efficiency of SPN Δ*ciaR* (response regulator) and Δ*ciaH* (its cognate histidine kinase) mutant strains in hBMECs were calculated as percentage survival at indicated time points relative to 0 h. (**B**) Comparison of acid tolerance response of SPN Δ*ciaH* and Δ*ciaH*Δ*spxB* mutant strains *in-vitro*, measured as percentage viability after exposure to lethal pH 4.4. (**C-D**) Percentage GFP positive population (**C**) and mean fluorescence intensity (**D**) of SPN:*P_htrA_*-GFP variants: WT, Δ*ciaH* and Δ*ciaHΔspxB*, grown in THY to 0.4 OD_600 nm_, analyzed by flow cytometry and expressed as percentage relative to WT:*P_htrA_*-GFP. (**E**) Comparison of intracellular survival efficiency of SPN Δ*ciaH* and Δ*ciaHΔspxB* mutant strains in hBMECs. (**F-H**) Comparison of acid tolerance response of WT and Δ*spxB* mutant strains in R6 (unencapsulated, serotype 2) (**F**) D39 (encapsulated, serotype 2) (**G**) and TIGR4 (encapsulated, serotype 4) (**H**) background *in-vitro*. Survival was measured as percentage viability after exposure to lethal pH 4.4. Data information: Experiments are performed thrice and data of representative experiments are presented as mean ± SD of triplicate wells. Statistical analysis was performed using Student’s two-tailed unpaired t-test (A-B, E-H) and one-way ANOVA with Tukey’s multiple comparison test (C, D). ns, non-significant; \**p*<0.05; \*\**p*<0.01; \*\*\**p*<0.001.

Overall, these confirm that SpxB contributes to acid stress tolerance, via regulation of CiaR and (in turn) its regulon. This was also reflected in the intracellular survival capability of these strains, with the Δc*iaH*Δ s*pxB* double mutant demonstrating reduced viability than the Δ*ciaH* mutant (~ 1.8-fold at 4 h.p.i and ~ 5.7-fold at 8 h.p.i) (Fig 2E). We also performed the *in vitro* ATR assay using WT and Δ*spxB* mutant strains and observed a drastic reduction in viability of the latter (~ 22-fold compared to WT) (Fig 2F). Since the 2 strains did not exhibit a difference in their normal growth rates (S2A Fig), it was confirmed that the difference in colony forming units (CFU) retrieved after exposure to lethal pH was indeed due to their differential viability under those conditions. This phenotype of reduced viability of Δ*spxB* mutant following exposure to lethal pH was also reproduced in the encapsulated strains D39 (serotype 2) and a serotype 4 strain TIGR4 (Fig 2G, H).

### Pyruvate oxidase is critical for pneumococcal survival inside the acidified vacuoles of brain endothelium for efficient transcytosis

We further explored the significance of SpxB in pneumococcal interaction with the brain endothelial cells. A Δ*spxB* mutant demonstrated reduced ability to adhere to and invade these cells, compared to WT (~ 1.7-fold in adherence and ~ 1.9 for invasion) (Fig 3A, B). Since Δ*spxB* mutant exhibited attenuated survival under acid stress *in vitro*, we performed intracellular survival assay and observed a similarly diminished ability of the mutant to survive inside the hBMECs (difference in ~ 3.8-fold at 4 h.p.i and ~ 12.4-fold at 6 h.p.i) (Fig 3C). By performing MTT cell viability assay, we confirmed that the difference in bacterial counts observed is not due to difference in viability of the cells infected with different strains (S2B Fig). To confirm whether the survival advantage conferred by SpxB is within the endocytic vacuoles of the brain endothelium, we created a Δ*ply* mutant strain. Pneumolysin (Ply), a pore forming toxin belonging to the family of cholesterol dependent cytolysins (CDCs) has been shown to damage the PCV membrane, triggering xenophagy [13]. Excessive damage to the vacuolar membrane allow pneumococci to escape into the cytosol where it would be recaptured by xenophagy or targeted to proteasomes, both of which has been implicated as host-protective mechanisms for the degradation of pneumococci invading the brain endothelium [13, 27]. Thus, a Δ*ply* mutant is forced to reside within intact endocytic vacuoles and demonstrate improved intracellular survival compared to WT [13]. When *spxB* was knocked out in the Δ*ply* background, the resulting double mutant Δ*ply*Δ*spxB* was severely attenuated for its intracellular survival capability confirming that SpxB is critical for SPN survival within the endocytic vacuoles of hBMECs (Fig 3D). We compared co-localization of WT and Δ*spxB* mutant strains with Lysotracker, at 6 h.p.i and observed no significant difference between the two, suggesting that the vacuoles harbouring these strains are acidified to similar extent (Fig 3E). However, the stark difference in viable count obtained for WT and Δ*spxB* mutant in the intracellular survival assays, at 4 h.p.i (which corresponds to 6 h.p.i. of the immunofluorescence studies) confirms that SpxB plays a role in improving pneumococcal viability within the acidified vacuoles, via activation of ATR.

**Fig 3:**
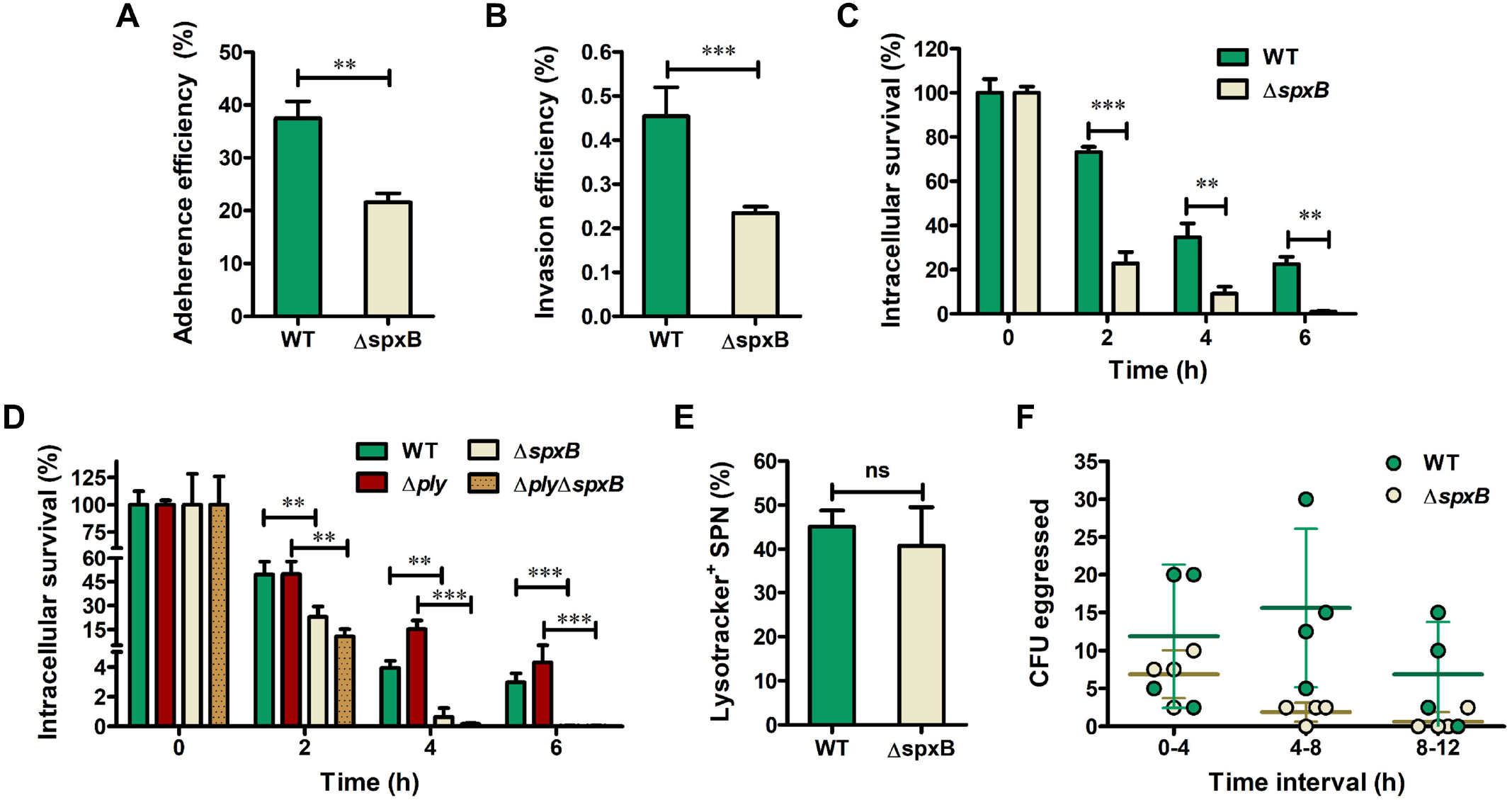
Pyruvate oxidase is critical for pneumococcal survival inside the acidified vacuoles of brain endothelium for effecient transcytosis. (**A-B**) Interaction of SPN WT and Δ*spxB* mutant strains with hBMECs assessed by adherence assay (**A**) and invasion assay (**B**). (**C**) Intracellular survival efficiency of SPN WT and Δ*spxB* mutant strains in hBMECs were calculated as percentage survival at indicated time points relative to 0 h. (**D**) Comparison of intracellular survival efficiency of SPN WT, Δ*ply, ΔspxB* and Δ*plyΔspxB* mutant strains in hBMECs. (**E**) Percentage co-localization of SPN WT and Δ*spxB* mutant strains with Lysotracker at 6 h.p.i. (**F**) Ability of SPN WT and Δ*spxB* mutant strains to egress out of hBMECs shown as CFU retrieved from cell supernatant during indicated time intervals. Data information: Experiments are performed thrice and data of representative experiments are presented as mean ± SD of triplicate wells. Statistical analysis was performed using Student’s two-tailed unpaired t-test (A-C, E) and one-way ANOVA with Tukey’s multiple comparison test (D). ns, non-significant; \**p*<0.05; \*\**p*<0.01; \*\*\**p*<0.001.

Previous work from our lab has demonstrated that, within an isogenic population of SPN, a subset which express low amount of Ply (SPN:Ply-Low) are predominantly vacuolar and demonstrate improved intracellular survival and transcytosis ability across the brain endothelium [27]. Intriguingly, deletion of *spxB* in SPN:Ply-Low background (SPN:Ply-Low Δ*spxB*) reduced its intracellular survival potential, further substantiating the role of SpxB in promoting pneumococcal survival inside the vacuoles (S2C Fig). Finally, we performed egression assay to assess the ability of SPN to transit out of BMECs. Δ*spxB* mutant strain exhibited a reduced trend in its ability to egress out of hBMECs (compared to WT), in corroboration with their reduced ability to survive inside the cells (Fig 3F). Overall, these results demonstrate that SpxB plays a critical role in pneumococcal survival inside BMECs, by enabling it to mount the ATR via CiaR regulation, to maintain viability in the acidic pH of vacuoles. Regulation of CiaR activity by SpxB is likely mediated by AcP, serving as an alternate phosphate donor and facilitating control of genes under the CiaR regulon. We also explored whether this newly established mechanism of SpxB in promoting intracellular persistence could be extrapolated to other host cell types, such as macrophages. Intracellular survival assays performed with WT and Δ*spxB* mutant strains in THP-1 macrophages failed to show a difference, suggesting that SpxB conferred advantage might be insufficient to prevent pneumococcal degradation by professional phagocytes (S2D Fig). This could be ascribed to differential mechanisms for improved pathogen elimination by macrophages such as the extent of phagosomal acidification, use of serine proteases and anti-microbial peptides and involvement of a distinct LC3-associated phagocytosis (LAP) pathway, as discussed by Inomata *et al* [28].

### Hydrogen peroxide and other reactive oxygen species confer survival advantage to pneumococci within the vacuoles of brain endothelial cells

Apart from the lethal low pH, the lysosomal compartment, where SPN has been found during trafficking through brain endothelium (Fig 1E, F), is abundant in degradative enzymes. These are detrimental for pneumococcal viability and SPN must adopt strategies to persist in such harsh environment. SPN produces significant amount of reactive oxygen species (ROS) in the form of hydrogen peroxide (H_2_O_2_) as a by-product of SpxB catalysed reaction (S2E Fig) and have mechanisms to withstand high concentration of ROS [29]. Interestingly, H_2_O_2_ has been implicated in interfering with the vacuolar maturation and cargo degradation process in diverse ways [30–32]. Hence, we next sought to investigate whether H_2_O_2_ produced by SPN can govern the fate of PCVs within BMECs. For this, we first assessed whether H_2_O_2_ generated via action of pneumococcal SpxB is sensed by hBMECs, by transfecting the endothelial cells with a genetically encoded probe HyPer-3 [33]. The fluorescence of HyPer increase as a result of conformational change brought about by oxidation, and is specific for detection of H_2_O_2_. Treatment of HyPer-transfected hBMECs with increasing concentrations of H_2_O_2_ showed a progressive increase in the median fluorescence intensity (MFI) of cells when analysed by flow cytometry (S3A Fig). Following infection with SPN, HyPer-transfected hBMECs demonstrated an increase in the MFI which was significantly higher for those infected with WT SPN than with Δ*spxB* mutant strain (~ 1.9-fold), indicating that H_2_O_2_ produced by SPN is sensed by the host cells (Fig 4A).

**Fig 4.**
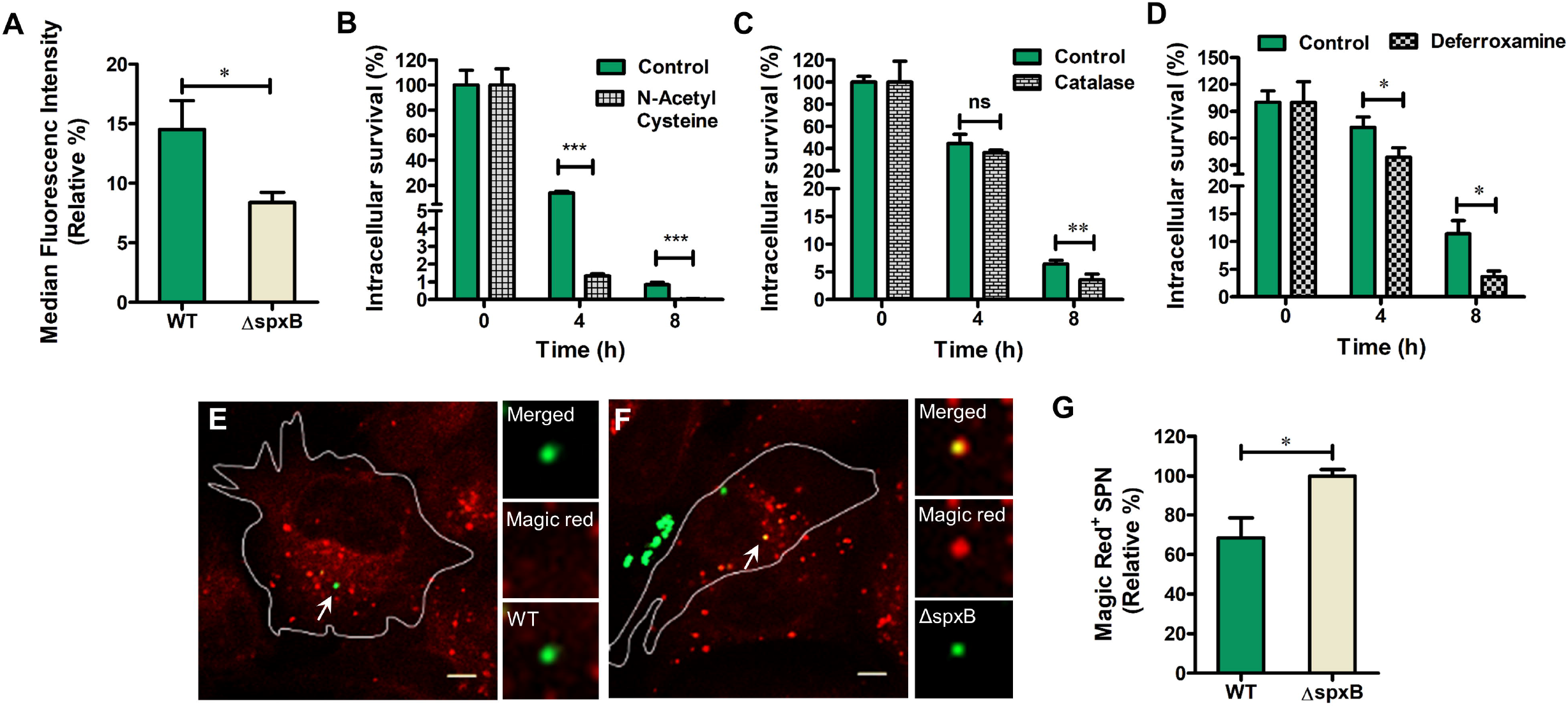
Hydrogen peroxide and other reactive oxygen species confer survival advantage to pneumococci within the vacuoles of brain endothelial cells. (**A**) Median fluorescence intensity of Hyper-transfected hBMECs infected with SPN WT and Δ*spxB* mutant strains, normalized to the fluorescence intensity of uninfected cells. (**B-D**) Intracellular survival efficiency of WT SPN in hBMECs treated with N-Acetyl Cysteine (**B**); 10 mM during 1 h pre-treatment and throughout the experiment, Catalase (**C**); 0.5 mg/ml during infection and 1 mg/ml in antibiotic containing medium and Deferoxamine (**D**); 50 μM during 3 h pre-treatment and 200 μM in antibiotic containing medium. (**E-F**) Confocal micrographs showing association of WT SPN (**E**) and Δ*spxB* (**F**) GFP variants (green) with Magic Red (red) at 6 h.p.i. Arrows designate the bacteria shown in insets. Positive events were localized at the following z-stack numbers: 6 out of 18 (for E) and 11 out of 21 (for F) (**G**). Percentage co-localization of WT SPN with Magic Red at 6 h.p.i, relative to that of Δ*spxB* mutant strain. Data information: Experiments are performed thrice and data of representative experiments are presented as mean ± SD of triplicate wells. Statistical analysis was performed using Student’s unpaired two-tailed t test (A-D, G). ns, non-significant; \**p*<0.05; \*\**p*<0.01; \*\*\**p*<0.001.

Next, in order to check whether this H_2_O_2_ influences the fate of PCVs, we treated hBMECs with different inhibitors of reactive oxygen species (ROS) (S3B Fig). N-acetyl cysteine (NAC) scavenges ROS directly owing to its thiol-nature [34] and also by increasing the levels of cellular glutathione (L-cysteine is a precursor of glutathione) [35]. Treatment of hBMECs with NAC and catalase which detoxifies H_2_O_2_ to water, reduced the intracellular survival potential of WT but not that of Δ*spxB* mutant strain (Fig 4B, C and S3C Fig). H_2_O_2_ is also converted to hydroxyl free radical via the Fenton reaction and the lysosomal compartment, owing to its rich iron content and low pH is presumed to favour this reaction [36]. Since hydroxyl radicals would be contained within the vacuoles owing to its charged nature (unlike H_2_O_2_) for a more localized effect and is a more potent ROS than H_2_O_2_, we checked if the hydroxyl radical formed from H_2_O_2_ could also be a contributor to pneumococcal persistence within BMEC vacuoles. For this, we treated the cells with deferoxamine, an iron chelator that would restrict the Fenton reaction and subsequent generation of hydroxyl radicals and observed that the survival of WT SPN within hBMECs was indeed reduced (Fig 4D). Work by Rybicka *et al* had previously demonstrated that H_2_O_2_ can reduce the proteolytic activity of lysosomes via oxidative inactivation of lysosomal cysteine cathepsins [32]. Based on this, we tested whether pneumococcal H_2_O_2_ could similarly interfere with the activity of lysosomal enzymes, by measuring the co-localization of WT and Δ*spxB* mutant strains with Magic Red Cathepsin B, a dye that fluoresce upon proteolytic cleavage by cathepsin B. Intriguingly, the degree of association of WT SPN with Magic Red, was significantly lower than that of Δ*spxB* mutant strain (Fig 4E-G and S3D Fig) implying deactivation of lysosomal cathepsin B by WT SPN. Taken together, these results indicate an important contribution of H_2_O_2_ and other ROS in prolonging pneumococcal intracellular survival, possibly by interfering with the activity of lysosomal hydrolases.

## DISCUSSION

The intricacies in the design of the endocytic pathway which equips a eukaryotic cell to perform diverse functions ranging from nutrient acquisition, signalling, recycling of plasma membrane components and defence against microbes, ironically offers multiple platforms for manipulation by pathogens to avert their intracellular degradation. *Listeria monocytogenes* and *S. pyogenes* have been known to use their pore forming toxins to damage the pathogen containing vacuole and escape into the cytosol [37, 38]. On the other hand, many others modify the identity of the vacuoles in which they reside to arrest its maturation and fusion with lysosomes. Common targets for manipulation by the pathogens such as *M. tuberculosis, S. typhimurium, L. pneumophila* etc. are phosphoinositides and Rab GTPases owing to their role in conferring identity to the endocytic compartments, which in turn governs its maturation [39–42]. *S. aureus* and *Helicobacter pylori* on the contrary, have been known to neutralize the acidity of their vacuoles by producing ammonia via action of urease [43, 44]. There are also rare examples such as *Coxiella brunetti* which in fact prefer acidic, lysosome-like compartment for their replication [45]. Such a lifestyle is facilitated by expression of factors that enable them to withstand the inhospitable phago-lysosomal compartment. For example, *S. aurues* modifies its cell wall to resist action of lysozyme [46]; staphylokinase and aureolysin helps in neutralizing alpha-defensins and the antimicrobial peptide LL-37 [47–49]. It also possess an arsenal of antioxidant enzymes that detoxify reactive oxygen and nitrogen species inside the phagolysosomes [50–53].

Although majority of the above examples refer to intracellular pathogens that reside within host cells for extensive period of time, evasion of endo-lysosomal degradation is a prerequisite even for those that are only transiently exposed to the intracellular milieu. This is the case for pneumococci that traverse the brain endothelial cells for invasion of the central nervous system. BMECs form the primary structural and functional constituent of the BBB and are characterized by the presence of tight junctions for strictly regulating the movement of cargoes across the barrier. This high trans-endothelial electrical resistance impedes transit through intercellular space and probably makes transcytosis a preferred route of BBB trafficking for pneumococci [10]. Previous studies have elucidated the role of xenophagy and ubiquitin-proteasome machineries in degradation of SPN invading the BMECs [13, 27]. These pathways were triggered upon damage to the PCVs brought about by pore forming activity of Ply. A Ply knockout mutant and pneumococcal subsets expressing low amounts of Ply were expectedly found to evade these pathways [13], confining their residence within intact vacuoles. Interestingly, although the default fate of endocytic vacuoles is lysosomal fusion, these strains demonstrated improved survival and transcytosis ability compared to WT [27]. We therefore speculated that SPN might have strategies to prevent their degradation in the endocytic compartments, possibly by interfering with its maturation. However, contrary to our expectations, we observed that majority of PCVs do undergo maturation and luminal acidification, with a subset even acquiring markers of lysosomes. This suggested that pneumococcal survival strategies within BMECs might be directed at withstanding the acidic pH and other degradative components of these lysosome-like vacuoles.

SPN possess an arsenal of 13 TCSs and an orphan response regulator which enable it to fine tune the gene expression to adapt to its environment [54]. Among these, CiaRH is the first and one of the most well characterized pneumococcal TCSs. Although the nature of the signal sensed by CiaH remains unknown, this TCS has been implicated in damage control response to lysis-inducing conditions, temperature, oxidative, and more relevant to our study, acid stress [23, 55–57]. Upon phosphorylation of aspartate residue at 51^st^ position (D51), CiaR binds to direct hexamer repeat TTTAAG and controls 15 promoters which regulates the expression of 24 genes [26]. Notable genes under the CiaR regulon include HtrA and PpmA (both implicated in virulence), those involved in teichoic acid synthesis (*lic1* operon) etc.

Significance of CiaRH in pneumococcal stress response is primarily attributed to its regulation of HtrA (high temperature responsive A) [58], which preserves pneumococcal cell integrity by acting as a chaperone (at low temperature) and a protease (at high temperature) to degrade proteins that are damaged beyond repair [59]. Consistent with the previous report on the role of CiaR in induction of pneumococcal ATR *in vitro* and within A549 epithelial cells [23], our results demonstrated a critical need for CiaR activation to preserve pneumococcal viability inside BMECs. Although the CiaR regulated genes responsible for mediating ATR requires further investigation, HtrA remains the primary candidate; we did observe activity of *htrA* promoter under lethal acidic stress. Additionally, many of the other CiaR regulated genes are also surface localized [26] suggesting their potential contribution in intra-vacuolar stress tolerance of SPN.

An earlier study had identified CiaRH transcription to be upregulated in response to host sialic acids [57]. Since pyruvate was one of the products of sialic acid metabolism, it was speculated that SpxB-mediated conversion to AcP may contribute to CiaR phosphorylation, in turn driving expression of genes involved in virulence [57]. High levels of expression of CiaR regulated genes even in the absence of CiaH favoured this theory where phosphorylation (and activation) of CiaR is mediated by another molecule [60]. Indeed, a recent study confirmed AcP-mediated phosphorylation of CiaR in SPN, independent of CiaH [24]. This process was found to be under strict (negative) regulation by acetate kinase (AckA) to prevent over activation of CiaR, which is lethal [24]. Our findings corroborate the contribution of SpxB in activation of CiaR, in turn, indirectly controlling the CiaR regulon. This role of SpxB was found to be essential for pneumococcal survival under lethal acidic stress *in vitro* as well as within the acidified vacuoles of BMECs, facilitating successful transcytosis.

Apart from CiaR, other factors have been implicated in pneumococcal response to acid stress. Probably the most important among these is the FoF1 ATPase (encoded by *atpABC*) which exclude protons from the bacterial cytoplasm and has been found to be essential for pneumococcal survival in RAW264.7 macrophages [23]. Analysis of transcriptional response in SPN following exposure to pH 6.0 revealed upregulation of genes involved in protein fate such as GrpE, DnaK, DnaJ chaperons and the ClpL, PrtA proteases. Additionally, there was also a trend towards increased glycolysis, evident by upregulation of the rate-limiting enzyme 1-phosphofructokinase and feeding of galactose intermediates into this pathway [21]. We believe that a combined action of CiaR (aided by SpxB) with these other mechanisms would preserve pneumococcal viability inside the acidified vacuoles during BBB trafficking. Intriguingly, increased glycolysis as a response to acid stress would lead to higher amount of pyruvate, which will aid in the production of more AcP via action of SpxB, improving pneumococcal viability under these conditions. Also upregulated in the transcriptional analysis study, were genes involved in oxidative stress response such as *gor* (glutathione reductase), subunits of Mn transporter (which is a cofactor for superoxide dismutase) and the *hslO* encoded chaperone [21]. We presume that these might assist pneumococci in protecting itself from the H_2_O_2_ generated as a by-product of SpxB activity. Since the contribution of SpxB in acid stress tolerance was significant not only in Δ*ciaH* knockout background, but even in the WT, there exists a possibility that SpxB might be operating in additional ways, distinct from CiaR activation, especially given the role of AcP as an intracellular messenger.

A noteworthy characteristic of pneumococci is its ability to generate large amounts of H_2_O_2_. Although SpxB serve as the primary source, pneumococcal H_2_O_2_ is also generated as a by-product of glycerol metabolism by the action of α–glycerophosphate oxidase (GlpO) [61]. ROS, including H_2_O_2_, has generally been observed as a host defence mechanism against pathogenic microbes, especially within the vacuoles of professional phagocytes [62]. A contrasting facet is the ability of ROS to manipulate the endo-lysosomal pathway in diverse ways, a feature, we hypothesized could be hijacked by SPN for improved survival within the vacuoles, given its ability to produce and withstand large amounts of H_2_O_2_. Survival assays performed by treating BMECs with ROS inhibitors: NAC and catalase, reduced the intracellular viable SPN count, supporting this theory. Treatment with deferoxamine, which would restrict the Fenton reaction and generation of hydroxyl radical, also showed a similar phenotype, confirming a supportive role of ROS in pneumococcal survival inside BMECs. H_2_O_2_ has been shown to deplete PI3P on early endosomes in a p38 MAPK-dependent manner to prevent endosomal maturation [30]. It is also capable of causing lysosomal membrane permeablization, leading to neutralization of luminal pH [31]. However, our immunofluorescence studies showed a significant association of WT SPN with the PI3P probe P40PX as well as with Lysotracker, suggesting a different mechanism for H_2_O_2_-aided pneumococcal survival. Another study demonstrated that H_2_O_2_ produced by eukaryotic NADPH oxidase (Nox2) inside the phagosomes of dendritic cells can reduce the proteolytic activity of lysosomes by oxidative inactivation of lysosomal cysteine cathepsins [32]. Cysteine cathpesins (B, C, F, H, K, L, O, S, V, X, W) require a reducing environment for their proteolytic activity and along with aspartic cathepsins (D, E), constitute the most abundant class of lysosomal proteases [63]. Our results of SPN co-localization with Magic Red Cathepsin B dye indicate that ROS generated via action of SpxB, within the PCVs can compromise the activity of lysosomal enzymes, specifically the cysteine cathepsins, enabling SPN to evade degradation within lysosomes for safe transcytosis.

Collectively, our studies suggest a dual role of SpxB in improving pneumococcal viability during transcytosis across BMECs (Fig 5). It would be interesting to explore whether SpxB contributes to pneumococcal survival in the recently discovered context of prolonged persistence and replication within splenic macrophages and cardiomyocytes [4, 5]. Owing to its role in generation of AcP and H_2_O_2_, SpxB has previously been implicated in pneumococcal pathogenesis, with SpxB mutant strains demonstrating attenuated virulence in animal models of colonization, pneumonia and sepsis [25]. AcP has been shown to modulate pneumococcal adherence to host cells [25] and regulate capsular expression [64]. It also act as a high-energy source of ATP, a feature which improves pneumococcal viability during oxidative stress [65]. H_2_O_2_, on the other hand, provides selective advantage to SPN during nasopharyngeal colonization, by inhibiting the growth of other microflora [66]. H_2_O_2_ has also been implicated as a cause of host genotoxicity and cytotoxicity [67, 68]. Peroxynitrates generated as a result of interaction of H_2_O_2_ with host derived nitric oxide and hydroxyl radicals generated via Fenton reaction exacerbates the lethality associate with pneumococcal infections [69]. Our studies provide additional insights into the significance of SpxB in pneumococcal virulence and its contribution in the pathogenesis of pneumococcal meningitis. Interestingly, diabetic patients have been linked with higher risk of invasive pneumococcal diseases (including meningitis) and have higher associated fatality rates, compared to non-diabetic people [70–72]. Although this correlation has mostly been thought to be a consequence of diabetes-related host infirmities [73], it is possible that increased blood glucose serving as a source of pyruvate and feeding into SpxB pathway, resulting in the generation of virulence mediators AcP and H_2_O_2_ could be aggravating pneumococcal disease burden in these patients. Therefore, SpxB, which is ubiquitously expressed by all SPN strains, could serve as a promising candidate for therapeutic interventions, providing options for better management of pneumococcal disease burden, even in elderly population.

**Fig 5.**
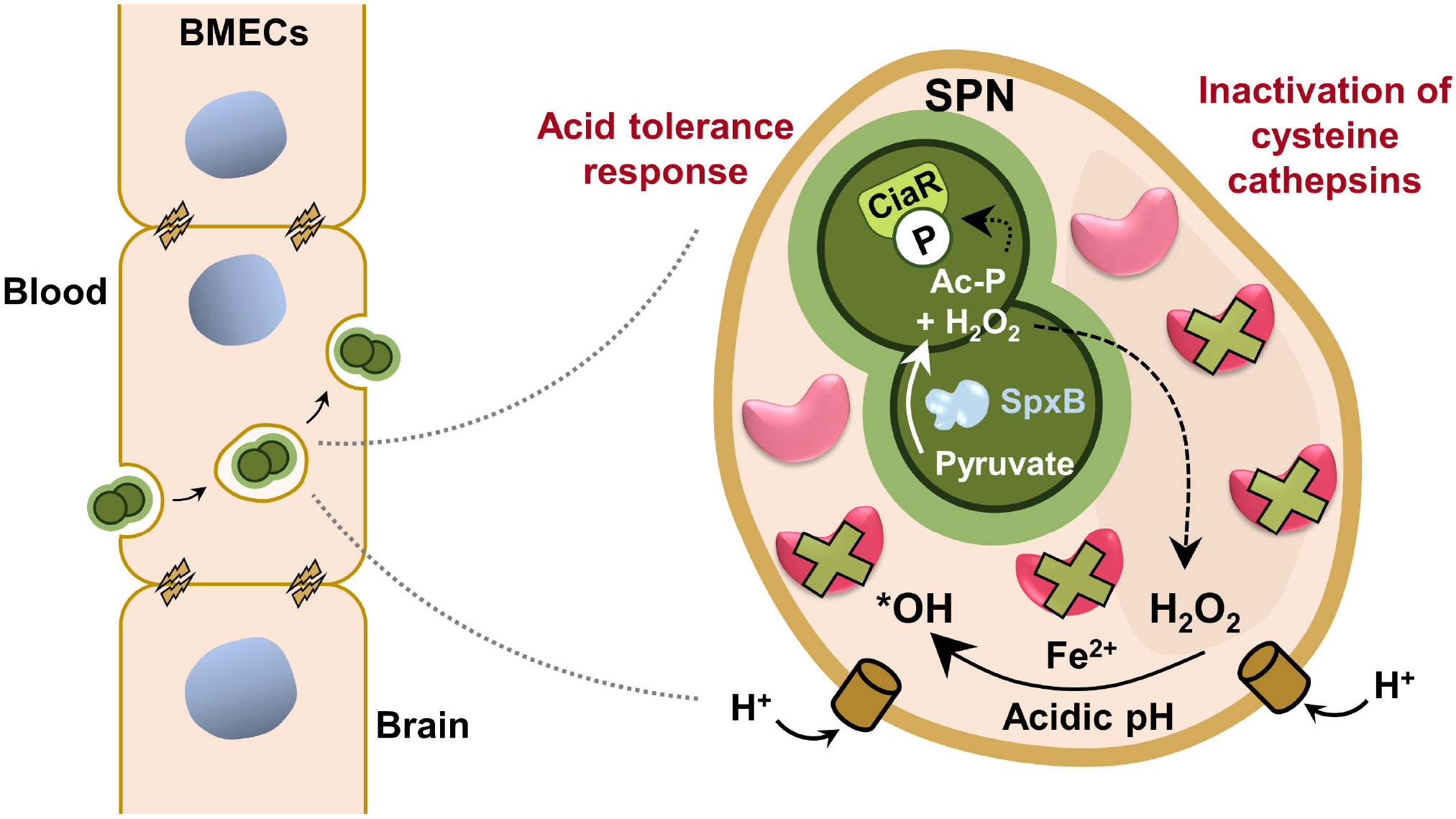
Schematic depicting contribution of SpxB in pneumococcal survival inside host cell endocytic vacuoles. SpxB, owing to its role in generation of AcP and H_2_O_2_, contributes to pneumococcal survival within the mature, acidified, lysosome-like vacuoles of brain endothelial cells. AcP contributes to induction of acid tolerance response, via phosphorylation and activation of CiaR, preserving pneumococcal viability under lethal acidic pH of the vacuoles. H_2_O_2_ (and hydroxyl radical formed via the fenton reaction), on the other hand can oxidize and inactivate lysosomal cysteine cathepsins, compromising the proteolytic/degradative capacity of lysosomes. These strategies facilitate successful transcytosis of pneumococci across host cellular barriers, including the blood-brain barrier.

## MATERIALS AND METHODS

### Cell culture

Human brain microvascular endothelial cells (hBMECs) were routinely cultured in RPMI 1640 (Gibco) supplemented with 10% fetal bovine serum (FBS, Gibco), 10% Nu-Serum (Corning) and 1% minimum essential medium non-essential amino acids (MEM-NEA, Gibco), on rat-tail collagen I (Corning) coated flasks/plates/coverslips at 37°C and 5% CO_2_. This cell line have been used as an *in vitro* BBB model for several host-pathogen interaction studies [74].

For the generation of P40PX-EGFP hBMECs, the cells were transfected (as described below) with a mammalian expression vector pAB121. This vector was constructed by sub-cloning the region encoding P40PX-EGFP (amplified from Addgene plasmid #19010) [19] into pMRX-IRES-Blast vector, kindly provided by Dr. T Yoshimori (Osaka University, Japan). The transfected cell line (hBMEC: P40PX-EGFP) were routinely grown in the presence of blasticidin (2 μg/ml).

THP-1 monocytes, kindly provided by Prof. S Mehra (IIT Bombay, India) were routinely cultured in RPMI 1640 supplemented with 10% FBS at 37°C and 5% CO_2_ as suspension culture. Differentiation of THP-1s into macrophages was achieved by treatment with 25 ng/ml phorbol 12-myristate13-acetate (PMA, Sigma-Aldrich) for 24 h followed by resting in fresh media for another 24 h.

### *Streptococcus pneumoniae* strains

*Streptococcus pneumoniae* strains D39 (encapsulated, serotype 2), R6 (non-encapsulated derivative of D39) and TIGR4 (encapsulated, serotype 4) were routinely grown in Todd Hewitt broth supplemented with 1.5% yeast extract (THY) or Brain Heart Infusion (BHI) agar at 37□C and 5% CO_2_. R6 and its derivatives were used for most assays (unless otherwise mentioned) as they exhibit improved adherence and invasion compared to their encapsulated counterparts [75], enabling better tracking of their intracellular fate. Although historically, non-encapsulated SPN (NESp) has been considered avirulent, recent reports indicate an increase in their association with invasive and non-invasive pneumococcal diseases [76, 77]; presumably, an outcome of selection pressure exerted by pneumococcal polysaccharide and conjugate vaccines which target encapsulated strains [78]. Pathogenesis of NESp is attributed to additional virulence mechanisms they possess, in order to compensate for the lack of capsule [76, 77, 79].

For the construction of Δ*spxB* mutant, the strategy of insertional inactivation using antibiotic resistance cassette was adopted. Briefly, *spxB* gene region containing the ORF (amplified from R6 genome using primer set 1-2) was cloned into pGEM-T Easy vector (Promega). *spxB* ORF was disrupted by cloning a kanamycin resistance cassette (amplified from pIBD38, kindly provided by I Biswas, KUMC, USA, using primer set 25-26) between the HpaI-BamHI restriction sites. The linearized plasmid (pAB103) was then used for the transformation of SPN using competence stimulating peptide (GenPro Biotech; CSP 1 for R6 / D39 and CSP 2 for TIGR4). The recombinants were selected using kanamycin (200 μg/ml) and gene disruption was confirmed by PCR using primers flanking the *spxB* gene (primer set 3-4). Functional loss of SpxB activity was confirmed by hydrogen peroxide assay. Same strategy was adopted for the generation of *spxB* mutants in the background of Δ*ciaH*, Δ*ply* [80] and Ply-Low [13] strains.

For the construction of R6Δ*ciaR* mutant, a cassette consisting of *ciaR* upstream and downstream regions flanking a spectinomycin resistance cassette (amplified from pUCSpec, kindly provided by I Biswas, KUMC, USA) was assembled in pBSK vector. *ciaR* and *ciaH* is an operon with a common promoter upstream of *ciaR* and the start codon of *ciaH* falls within the *ciaR* ORF. Hence, to construct a *ciaR* deletion mutant without affecting expression of *ciaH*, the cassette was constructed in such a way that the promoter and other regulatory elements upstream of *ciaR* (in the WT genome) were shifted downstream of the antibiotic resistance cassette and immediately upstream of *ciaH* ORF. Briefly, the downstream region consisting of the 5’ untranslated region (UTR) of *ciaR* (primer set 7-8) fused to *ciaH* ORF (primer set 9-10) was generated by overlap extension PCR and cloned into BamHI and XbaI sites of pBSK. Next, the *ciaR* upstream region (primer set 5-6), was cloned between XhoI and EcoRI sites. The spectinomycin resistance cassette (primer set 29-30) was finally cloned between EcoRI and BamHI sites. The resultant construct (pAB147) was used to transform SPN strain R6 to generate a Δ*ciaR* mutant after selection on spectinomycin (100 μg/ml). *R6*Δ*ciaH* was similarly constructed using a cassette consisting of *ciaH* upstream region (primer set 11-12), spectinomycin resistance cassette and *ciaH* downstream region (primer set 13-14) (pAB153). Restriction sites used for cloning the inserts were same as above. For deletion of complete *ciaRH* operon, a cassette consisting of *ciaR* upstream region, chloramphenicol resistance cassette (amplified from pPEP1, Addgene plasmid #61046 [81], using primer set 27-28) and *ciaH* downstream region was constructed (pAB168) and used for transformation of SPN R6. Recombinants were selected on chloramphenicol (2 μg/ml). For the construction of R6:CiaR^D51A^, *ciaRH* operon (primer set 15-16) was first cloned into pBSK between EcoRI-BamHI sites and used as a template for the generation of CiaR^D51A^ site directed mutation using primer set 17-18 by inverse PCR. Next a cassette consisting of *ciaR* upstream region, spectinomycin resistance cassette, *ciaRH* operon containing the CiaR^D51A^ mutation and *ciaH* downstream region was assembled in pBSK (pAB178) and used for the transformation of *R6*Δ*ciaRH* generated earlier. The recombinants were selected on spectinomycin and the mutation was confirmed by DNA sequencing. Proper recombination in all the *ciaRH* variants were verified by PCR of the *ciaRH* gene loci using primers flanking the region (primer set 19-20).

For construction of R6:*P_htrA_*-GFP strain, the 5’UTR of *htrA* (amplified from R6 genome, using primer set 21-22) was cloned upstream of GFP ORF in the plasmid pAB309. The *P_htrA_*-GFP cassette was then sub-cloned into the XbaI-XhoI restriction sites in pPEP1 vector (Addgene plasmid #61046) and used for the transformation of R6. Recombinants were selected on spectinomycin and confirmed by PCR using flanking primers (primer set 23-24). GFP expression was confirmed by fluorescence microscopy and flow cytometry analysis. For generation of R6:*P_htrA_*-GFPΔ*ciaH*, R6:*P_htrA_*-GFP strain was transformed with a *ciaH* deletion construct pAB167 (same as pAB153, but harboring a chloramphenicol resistance cassette). R6:*P_htrA_*-GFPΔ*ciaH*Δ*spxB* was generated by transforming R6:*P_htrA_*-GFPΔ*ciaH* with linearized pAB103.

SPN GFP and RFP fluorescent variants for immunofluorescence microscopy studies were generated by transformation of the respective strains with *hlpA*-GFP/tagRFP cassette (kindly provided by J-W Veening, University of Lausanne, Switzerland) [82] and selected using chloramphenicol (4.5 μg/ml), followed by confirmation with PCR using the flanking primer set 33-34 and fluorescence microscopy.

A list of all the SPN strains used in this study and primers used in the construction of these strains are summarized in S1 Table and S2 Table, respectively.

### Antibodies and reagents

Primary antibodies used for immunofluorescence microscopy were anti-phosphoryl choline (Sigma-Aldrich, M1421) and anti-cathepsin B (Abcam, ab58802). Anti-serum against SPN enolase was a gift from S Hammerschmidt (University of Greifswald, Germany). Lysotracker Deep Red (Invitrogen, L12492), a fluorescent acidotropic dye was used to label acidified vacuoles: late endosomes and lysosomes. Magic Red Cathepsin B Assay (ImmunoChemistry Technologies, #937) was used as an indicator for active cysteine cathepsins. Phorbol 12-myristate 13-acetate (PMA, P8139), N-acetyl cysteine (NAC, A9165), catalase (C9322) and deferoxamine (D9533) were procured from Sigma.

### Hydrogen peroxide assay

The ability of SPN WT and mutant strains to produce H_2_O_2_ was measured by a biochemical assay [65]. Briefly, SPN cultures grown in THY, to mid-exponential phase (OD_600 nm_ 0.4) were pelleted and OD_600 nm_ adjusted to 0.4 in PBS + 1% glucose. 180 μl of this sample (and its dilutions) was mixed with a 20 μl solution consisting of 3 mg/ml of 2,2’-azinobis-(3-ethylbenzthiazolinesulfonic acid) (ABTS) and 0.2 mg/ml of horseradish peroxidase (HRP) in 0.1 M sodium phosphate buffer (pH 7.0), in a 96-well flat bottom plate. The reaction mixture was incubated at 37°C for 30 min. The plate was centrifuged at 1500 g for 3 min and the supernatants were transferred to fresh wells. The absorbance was measured at 560 nm using a microplate spectrophotometer (Thermo Fischer Scientific). Amount of H_2_O_2_ produced was calculated by comparison to a standard curve that was generated using known concentrations of H_2_O_2_ (Merck).

### Intracellular survival assay

hBMEC monolayers grown on 24-well plates were washed thrice with RPMI and incubated in assay medium (RPMI + 10% FBS) for 1 h. SPN cultures grown to OD_600 nm_ 0.4 were pelleted, washed with PBS and finally OD_600 nm_ adjusted to 0.4 in PBS (~10^8^ CFU/ml). After necessary dilution in assay media to achieve multiplicity of infection (MOI) 10, 100 μl of this inoculum was added to the monolayer of hBMECs. Dilutions of the inoculum were spotted on BHI agar to enumerate the exact inoculum count. The plate was centrifuged at 800 rpm for 5 min to synchronize the infection and incubated at 37°C and 5% CO_2_ for 1 h. Following infection, the cells were washed thrice with RPMI to remove unbound and loosely adhered SPN and incubated with fresh assay medium containing 10 μg/ml penicillin G and 400 μg/ml gentamicin for 2 h to kill extracellular SPN. After washing thrice with RPMI, the cells were detached by adding 100 μl of 0.05% trypsin (Gibco) and lysed with 400 μl of 0.025% Triton X-100. The lysate was then spread plated on BHI agar to enumerate SPN CFU. Invasion percentage was calculated as follows: (CFU of SPN in the lysate / CFU of SPN in the inoculum) * 100. For intracellular survival assays, at different time points post antibiotic treatment, the cells were lysed and spread plated as described earlier. Percentage of viable intracellular SPN at any time point was calculated as follows: (CFU of SPN in the lysate at X h / CFU of SPN in the lysate at 0 h) * 100. For intracellular survival assays in the presence of inhibitors, hBMECs were pre-treated with N-acetyl cysteine (10 mM, 1 h) or catalase (0.5 mg/ml, 1 h) or deferoxamine (50 μM, 3 h) and the treatment remained throughout the experiment. To determine the adherence efficiency, hBMECs were washed with RPMI 6 times after 30 min of infection with SPN and lysed as described earlier. Dilutions of the lysates were spotted on BHI agar to enumerate CFU of adherent SPN and was expressed as a percentage of the inoculum. THP-1 macrophages were infected with SPN at MOI of 0.1 for 20 min and antibiotic treatment was reduced to 1 h.

### Egression assay

Monolayer of hBMECs were infected with SPN strains at MOI 10 for 1 h. After 2 h of incubation in the antibiotic containing medium (10 μg/ml penicillin G and 400 μg/ml gentamicin), the cells were washed and incubated in fresh assay medium containing 0.04 μg/ml of penicillin G (bacteriostatic concentration). Every 4 h, this media was replaced and the old supernatant was spread plated on BHI agar to enumerate the CFU of SPN egressed / recycled out of the cell.

### MTT cell viability assay

Effect of SPN infection or inhibitor treatments on the viability of hBMECs were quantified using EZcount™ MTT Cell Assay Kit as per manufacturer’s protocol (HiMedia).

### Immunofluorescence microscopy

For immunofluorescence microscopy experiments, hBMECs were grown on collagen-coated glass coverslips in 24-well plates. Infection was performed as described earlier using MOI of 25. Either GFP/RFP-tagged SPN strains were used for infection or unlabelled SPN strains were stained with anti-phosphoryl choline or anti-enolase antibody post fixation. For Lysotracker Deep Red staining, the dye was added to the cell culture media at a final concentration of 75-100 nM, 1 h before fixation. Magic Red Cathepsin B staining was performed as per manufacturer’s protocol; the dye was added to the cell culture media 1 h before fixation. At desired time points post infection, the cells were fixed by adding 4% paraformaldehyde (PFA, Sigma-Aldrich) for 15 min at room temperature (RT) or chilled methanol for 10 min at −20°C (for cathepsin B staining). The cells were then permeabilized (for PFA fixed samples) using 0.1% Triton X-100 for 10 min at RT and blocked using 3% bovine serum albumin (BSA, Himedia) for 2 h at RT. Post blocking, primary antibody (diluted in 1% BSA) was added and incubated at RT for 2 h or 4°C overnight. After washing to remove unbound antibody, the samples were incubated with fluorophore-conjugated secondary antibody at RT for 1 h. To selectively label extracellular SPN, the antibody staining was performed prior to permeabilization. Finally, the coverslips were mounted on glass slides using Vectashield mounting media containing DAPI (Vector Laboratories).

The samples were imaged using oil immersion Plan-Apochromat 40X/1.3 NA or iPlan-Apochromat 63X/1.4 NA objective of a confocal laser scanning microscope (Carl Zeiss, LSM780). Images were acquired after optical sectioning and processed using ZEN lite software (Version 5.0). For quantitation, at least 100 intracellular bacteria were counted per coverslip in triplicates (unless mentioned otherwise).

### Acid tolerance response assay

SPN WT and mutant strains were grown in THY (pH 7.1) to OD_600nm_ 0.3. The cells were then pelleted and resuspended in THY, pH 5.9 (sub-lethal pH) and incubated for 2 h at 37°C. The culture was then diluted 1:10 in THY, pH 4.4 (lethal pH) and incubated for another 2 h at 37°C. Dilutions of the cultures were spotted on BHI agar before and after incubation in pH 4.4 to enumerate viable SPN. CFU post incubation was expressed as percentage of initial CFU to calculate percentage survival under lethal acidic stress [23].

### hBMEC transfection

Transfections were performed using Lipofectamine 2000 (Invitrogen) according to manufacturer’s protocol. Briefly, 0.5 μg of plasmid DNA (in Opti-MEM), isolated using Plasmid mini kit (Qiagen), was mixed with 1 μl Lipofectamine 2000 (in OptiMEM) and added to 80% confluent hBMECs. After 5 h of incubation at 37° C, the medium was replaced with fresh culture medium. For the generation of stable cell lines, 48 h following transfection, hBMECs were transferred to medium containing the desired antibiotic for 2-3 weeks, following which the antibiotic resistance colonies were pooled for further propagation.

For transient transfection of hBMECs with the plasmid pC1-HyPer-3, (Addgene plasmid #42131) [33], hBMEC monolayers were transfected as described above. After 24 h, the cells were used for SPN infection or H_2_O_2_ treatment and analysed by flow cytometry.

### Flow cytometry

24 h post transfection with pC1-HyPer-3, hBMECs were infected with SPN WT and Δ*spxB* mutant strains at MOI 10 for 1 h. The cells were harvested by trypsinization, washed and resuspended in PBS containing 0.5% BSA. For control experiments, H_2_O_2_ was added to the cell suspension 5 min before initiation of the flow. The 488-nm excitation laser and 530±15 nm emission filter of the BD FACS Aria flow cytometer was used. Data was analyzed using FlowJo software.

For flow cytometry analysis of R6:*P_htrA_*-GFP variants, the cells were incubated in THY media of different pH as described in the method for acid tolerance response assay and then fixed with 4% PFA. The cells were washed and resuspended in PBS and analyzed as described above.

### Statistical analysis

GraphPad Prism version 5 was used for statistical analysis. Statistical tests undertaken for individual experiments are mentioned in the respective figure legends. Data are presented as mean ± standard deviation (SD) unless mentioned otherwise. Statistical significance was accepted at *p*<0.05. \*\*\**p* <0.001, ** *p* <0.01, \**p* <0.05, ns-non significant.

## Supporting information

Relative contribution of CiaRH and pyruvate oxidase towards pneumococcal tolerance to acidic stress

Pyruvate oxidase modulates pneumococcal interaction with the brain endothelial cells

Contribution of hydrogen peroxide to pneumococcal survival within the vacuoles of brain endothelial cells

List of strains

List of primers

## ACKNOWLEDGEMENTS

We acknowledge the Biosafety Level 2 facility, Confocal Microscopy facility and Flow Cytometry facility at IIT Bombay. AA acknowledge financial support from CSIR, Govt. of India.

## SUPPLEMENTARY INFORMATION CAPTIONS

**S1 Fig. Relative contribution of CiaRH two-component system and pyruvate oxidase towards pneumococcal tolerance to acidic stress.**

(**A**) Intracellular survival efficiency of SPN WT, Δ*ciaR* and CiaR^D51A^ (non-phosphorylatable) mutant strains in hBMECs were calculated as percentage survival at indicated time points relative to 0 h. (**B-C**) Comparison of acid tolerance response of SPN WT, Δ*ciaRH* and Δ*ciaR* and CiaR^D51A^ mutant strains *in-vitro*, measured as percentage viability after exposure to lethal pH 4.4. (**D**) Intracellular survival efficiency of SPN WT, Δ*ciaR* and Δ*ciaH* mutant strains in hBMECs were calculated as percentage survival at indicated time points relative to 0 h. (**E**) Percentage GFP positive population of SPN:*P_htrA_*-GFP variants: WT, Δ*ciaH* and Δ*ciaH*Δ*spxB*, upon exposure to lethal acid stress (protocol in methods section), analyzed by flow cytometry. Percentage GFP positive population in WT:*P_htrA_*-GFP under acidic stress was considered as 100%.

Data information: Experiments are performed thrice and data of representative experiments are presented as mean ± SD of triplicate wells or as mean±SEM of 2 independent experiments (E). Statistical analysis was performed using one-way ANOVA with Tukey’s multiple comparison test (A-D). ns, non-significant; \**p*<0.05; \*\**p*<0.01; \*\*\**p*<0.001.

**S2 Fig. Pyruvate oxidase modulates pneumococcal interaction with the brain endothelial cells.** (**A**) Growth kinetics of SPN WT and Δ*spxB* mutant strains in THY. (**B**) hBMEC viability assay performed at 12 h.p.i with SPN WT and Δ*spxB* mutant strains. (**C**) Comparison of intracellular survival efficiency of SPN:Ply-Low and SPN:Ply-Low Δ*spxB* mutant strains in hBMECs. (**D**) Intracellular survival efficiency of SPN WT and Δ*spxB* mutant strains in THP-1 macrophages were calculated as percentage survival at indicated time points relative to 0 h. (**E**) Hydrogen peroxide assay to verify loss of SpxB activity in Δ*spxB* mutant strain.

Data information: Experiments are performed thrice and data of representative experiments are presented as mean ± SD of triplicate wells. Statistical analysis was performed using Student’s unpaired two-tailed t test (C-E) and one-way ANOVA with Tukey’s multiple comparison test (B). ns, non-significant; \**p*<0.05; \*\**p*<0.01; \*\*\**p*<0.001.

**S3 Fig. Contribution of hydrogen peroxide to pneumococcal survival within the vacuoles of brain endothelial cells.** (**A**) Median fluorescence intensity of Hyper-transfected hBMECs treated with increasing concentrations of H_2_O_2_. (**B**) Cell viability assay performed on hBMECs treated with reactive oxygen species inhibitors: N-Acetyl Cysteine, catalase or deferroxamine, at 12 h.p.i with WT SPN. (**C**) Intracellular survival efficiency of Δ*spxB* mutant strain in hBMECs following treatment with N-Acetyl Cysteine; 10 mM during 1 h pre-treatment and throughout the experiment (**D**) Percentage co-localization of SPN WT and Δ*spxB* mutant strains with cathepsin B at 6 h.p.i.

Data information: Experiments are performed thrice and data of representative experiments are presented as mean ± SD of triplicate wells. Statistical analysis was performed using Student’s unpaired two-tailed t test (C, D) and one-way ANOVA with Tukey’s multiple comparison test (B). ns, non-significant; \**p*<0.05; \*\**p*<0.01; \*\*\**p*<0.001.

**S1 Table. List of *S. pneumoniae* strains.**

**S2 Table. List of primers.**

## REFERENCES

1. Osterlund A, Popa R, Nikkila T, Scheynius A, Engstrand L. Intracellular reservoir of Streptococcus pyogenes in vivo: a possible explanation for recurrent pharyngotonsillitis. The Laryngoscope. 1997;107(5):640–7. doi: 10.1097/00005537-199705000-00016. PubMed PMID: 9149167.

2. O’Neill AM, Thurston TL, Holden DW. Cytosolic Replication of Group A Streptococcus in Human Macrophages. mBio. 2016;7(2):e00020–16. doi: 10.1128/mBio.00020-16. PubMed PMID: 27073088; PubMed Central PMCID: PMC4959517.

3. Rollin G, Tan X, Tros F, Dupuis M, Nassif X, Charbit A, et al. Intracellular Survival of Staphylococcus aureus in Endothelial Cells: A Matter of Growth or Persistence. Frontiers in microbiology. 2017;8:1354. doi: 10.3389/fmicb.2017.01354. PubMed PMID: 28769913; PubMed Central PMCID: PMC5515828.

4. Ercoli G, Fernandes VE, Chung WY, Wanford JJ, Thomson S, Bayliss CD, et al. Intracellular replication of Streptococcus pneumoniae inside splenic macrophages serves as a reservoir for septicaemia. Nature microbiology. 2018;3(5):600–10. doi: 10.1038/s41564-018-0147-1. PubMed PMID: 29662129; PubMed Central PMCID: PMC6207342.

5. Brissac T, Shenoy AT, Patterson LA, Orihuela CJ. Cell Invasion and Pyruvate Oxidase-Derived H2O2 Are Critical for Streptococcus pneumoniae-Mediated Cardiomyocyte Killing. Infection and immunity. 2018;86(1). doi: 10.1128/IAI.00569-17. PubMed PMID: 29061707; PubMed Central PMCID: PMC5736805.

6. Hyams C, Camberlein E, Cohen JM, Bax K, Brown JS. The Streptococcus pneumoniae capsule inhibits complement activity and neutrophil phagocytosis by multiple mechanisms. Infection and immunity. 2010;78(2):704–15. doi: 10.1128/IAI.00881-09. PubMed PMID: 19948837; PubMed Central PMCID: PMC2812187.

7. Andre GO, Converso TR, Politano WR, Ferraz LF, Ribeiro ML, Leite LC, et al. Role of Streptococcus pneumoniae Proteins in Evasion of Complement-Mediated Immunity. Frontiers in microbiology. 2017;8:224. doi: 10.3389/fmicb.2017.00224. PubMed PMID: 28265264; PubMed Central PMCID: PMC5316553.

8. Dalia AB, Standish AJ, Weiser JN. Three surface exoglycosidases from Streptococcus pneumoniae, NanA, BgaA, and StrH, promote resistance to opsonophagocytic killing by human neutrophils. Infection and immunity. 2010;78(5):2108–16. doi: 10.1128/IAI.01125-09. PubMed PMID: 20160017; PubMed Central PMCID: PMC2863504.

9. Janoff EN, Rubins JB, Fasching C, Charboneau D, Rahkola JT, Plaut AG, et al. Pneumococcal IgA1 protease subverts specific protection by human IgA1. Mucosal immunology. 2014;7(2):249–56. doi: 10.1038/mi.2013.41. PubMed PMID: 23820749; PubMed Central PMCID: PMC4456019.

10. Iovino F, Orihuela CJ, Moorlag HE, Molema G, Bijlsma JJ. Interactions between blood-borne Streptococcus pneumoniae and the blood-brain barrier preceding meningitis. PloS one. 2013;8(7):e68408. doi: 10.1371/journal.pone.0068408. PubMed PMID: 23874613; PubMed Central PMCID: PMC3713044.

11. Anil A, Banerjee A. Pneumococcal Encounter With the Blood-Brain Barrier Endothelium. Frontiers in cellular and infection microbiology. 2020;10:590682. doi: 10.3389/fcimb.2020.590682. PubMed PMID: 33224900; PubMed Central PMCID: PMC7669544.

12. Surve MV, Apte S, Bhutda S, Kamath KG, Kim KS, Banerjee A. Streptococcus pneumoniae utilizes a novel dynamin independent pathway for entry and persistence in brain endothelium. Current Research in Microbial Sciences. 2020.

13. Surve MV, Bhutda S, Datey A, Anil A, Rawat S, Pushpakaran A, et al. Heterogeneity in pneumolysin expression governs the fate of Streptococcus pneumoniae during blood-brain barrier trafficking. PLoS pathogens. 2018;14(7):e1007168. doi: 10.1371/journal.ppat.1007168. PubMed PMID: 30011336; PubMed Central PMCID: PMC6062133.

14. Iovino F, Hammarlof DL, Garriss G, Brovall S, Nannapaneni P, Henriques-Normark B. Pneumococcal meningitis is promoted by single cocci expressing pilus adhesin RrgA. The Journal of clinical investigation. 2016;126(8):2821–6. doi: 10.1172/JCI84705. PubMed PMID: 27348589; PubMed Central PMCID: PMC4966305.

15. Ring A, Weiser JN, Tuomanen EI. Pneumococcal trafficking across the blood-brain barrier. Molecular analysis of a novel bidirectional pathway. The Journal of clinical investigation. 1998;102(2):347–60. doi: 10.1172/JCI2406. PubMed PMID: 9664076; PubMed Central PMCID: PMC508893.

16. Huotari J, Helenius A. Endosome maturation. The EMBO journal. 2011;30(17):3481–500. doi: 10.1038/emboj.2011.286. PubMed PMID: 21878991; PubMed Central PMCID: PMC3181477.

17. Behnia R, Munro S. Organelle identity and the signposts for membrane traffic. Nature. 2005;438(7068):597–604. doi: 10.1038/nature04397. PubMed PMID: 16319879.

18. Schink KO, Raiborg C, Stenmark H. Phosphatidylinositol 3-phosphate, a lipid that regulates membrane dynamics, protein sorting and cell signalling. BioEssays: news and reviews in molecular, cellular and developmental biology. 2013;35(10):900–12. doi: 10.1002/bies.201300064. PubMed PMID: 23881848.

19. Kanai F, Liu H, Field SJ, Akbary H, Matsuo T, Brown GE, et al. The PX domains of p47phox and p40phox bind to lipid products of PI(3)K. Nature cell biology. 2001;3(7):675–8. doi: 10.1038/35083070. PubMed PMID: 11433300.

20. Turk B, Turk D, Turk V. Lysosomal cysteine proteases: more than scavengers. Biochimica et biophysica acta. 2000;1477(1-2):98–111. doi: 10.1016/s0167-4838(99)00263-0. PubMed PMID: 10708852.

21. Martin-Galiano AJ, Overweg K, Ferrandiz MJ, Reuter M, Wells JM, de la Campa AG. Transcriptional analysis of the acid tolerance response in Streptococcus pneumoniae. Microbiology. 2005;151(Pt 12):3935–46. doi: 10.1099/mic.0.28238-0. PubMed PMID: 16339938.

22. Halfmann A, Schnorpfeil A, Muller M, Marx P, Gunzler U, Hakenbeck R, et al. Activity of the two-component regulatory system CiaRH in Streptococcus pneumoniae R6. Journal of molecular microbiology and biotechnology. 2011;20(2):96–104. doi: 10.1159/000324893. PubMed PMID: 21422763.

23. Cortes PR, Pinas GE, Cian MB, Yandar N, Echenique J. Stress-triggered signaling affecting survival or suicide of Streptococcus pneumoniae. International journal of medical microbiology: IJMM. 2015;305(1):157–69. doi: 10.1016/j.ijmm.2014.12.002. PubMed PMID: 25543170.

24. Kaiser S, Hoppstadter LM, Bilici K, Heieck K, Bruckner R. Control of acetyl phosphate-dependent phosphorylation of the response regulator CiaR by acetate kinase in Streptococcus pneumoniae. Microbiology. 2020;166(4):411–21. doi: 10.1099/mic.0.000894. PubMed PMID: 32553069.

25. Spellerberg B, Cundell DR, Sandros J, Pearce BJ, Idanpaan-Heikkila I, Rosenow C, et al. Pyruvate oxidase, as a determinant of virulence in Streptococcus pneumoniae. Molecular microbiology. 1996;19(4):803–13. doi: 10.1046/j.1365-2958.1996.425954.x. PubMed PMID: 8820650.

26. Halfmann A, Kovacs M, Hakenbeck R, Bruckner R. Identification of the genes directly controlled by the response regulator CiaR in Streptococcus pneumoniae: five out of 15 promoters drive expression of small non-coding RNAs. Molecular microbiology. 2007;66(1):110–26. doi: 10.1111/j.1365-2958.2007.05900.x. PubMed PMID: 17725562.

27. Iovino F, Gradstedt H, Bijlsma JJ. The proteasome-ubiquitin system is required for efficient killing of intracellular Streptococcus pneumoniae by brain endothelial cells. mBio. 2014;5(4):e00984–14. doi: 10.1128/mBio.00984-14. PubMed PMID: 24987087; PubMed Central PMCID: PMC4161243.

28. Inomata M, Xu S, Chandra P, Meydani SN, Takemura G, Philips JA, et al. Macrophage LC3-associated phagocytosis is an immune defense against *Streptococcus pneumoniae* that diminishes with host aging. Proceedings of the National Academy of Sciences. 2020:202015368. doi: 10.1073/pnas.2015368117.

29. Yesilkaya H, Andisi VF, Andrew PW, Bijlsma JJ. Streptococcus pneumoniae and reactive oxygen species: an unusual approach to living with radicals. Trends in microbiology. 2013;21(4):187–95. doi: 10.1016/j.tim.2013.01.004. PubMed PMID: 23415028.

30. Kano F, Arai T, Matsuto M, Hayashi H, Sato M, Murata M. Hydrogen peroxide depletes phosphatidylinositol-3-phosphate from endosomes in a p38 MAPK-dependent manner and perturbs endocytosis. Biochimica et biophysica acta. 2011;1813(5):784–801. doi: 10.1016/j.bbamcr.2011.01.023. PubMed PMID: 21277337.

31. Denamur S, Tyteca D, Marchand-Brynaert J, Van Bambeke F, Tulkens PM, Courtoy PJ, et al. Role of oxidative stress in lysosomal membrane permeabilization and apoptosis induced by gentamicin, an aminoglycoside antibiotic. Free radical biology & medicine. 2011;51(9):1656–65. doi: 10.1016/j.freeradbiomed.2011.07.015. PubMed PMID: 21835240.

32. Rybicka JM, Balce DR, Chaudhuri S, Allan ER, Yates RM. Phagosomal proteolysis in dendritic cells is modulated by NADPH oxidase in a pH-independent manner. The EMBO journal. 2012;31(4):932–44. doi: 10.1038/emboj.2011.440. PubMed PMID: 22157818; PubMed Central PMCID: PMC3280544.

33. Bilan DS, Pase L, Joosen L, Gorokhovatsky AY, Ermakova YG, Gadella TW, et al. HyPer-3: a genetically encoded H(2)O(2) probe with improved performance for ratiometric and fluorescence lifetime imaging. ACS chemical biology. 2013;8(3):535–42. doi: 10.1021/cb300625g. PubMed PMID: 23256573.

34. Moldeus P, Cotgreave IA, Berggren M. Lung protection by a thiol-containing antioxidant: N-acetylcysteine. Respiration; international review of thoracic diseases. 1986;50 Suppl 1:31–42. doi: 10.1159/000195086. PubMed PMID: 3809741.

35. Meister A, Anderson ME. Glutathione. Annual review of biochemistry. 1983;52:711–60. doi: 10.1146/annurev.bi.52.070183.003431. PubMed PMID: 6137189.

36. Kurz T, Terman A, Gustafsson B, Brunk UT. Lysosomes in iron metabolism, ageing and apoptosis. Histochemistry and cell biology. 2008;129(4):389–406. doi: 10.1007/s00418-008-0394-y. PubMed PMID: 18259769; PubMed Central PMCID: PMC2668650.

37. Schnupf P, Portnoy DA. Listeriolysin O: a phagosome-specific lysin. Microbes and infection. 2007;9(10):1176–87. doi: 10.1016/j.micinf.2007.05.005. PubMed PMID: 17720603.

38. O’Seaghdha M, Wessels MR. Streptolysin O and its co-toxin NAD-glycohydrolase protect group A Streptococcus from Xenophagic killing. PLoS pathogens. 2013;9(6):e1003394. doi: 10.1371/journal.ppat.1003394. PubMed PMID: 23762025; PubMed Central PMCID: PMC3675196.

39. Vergne I, Chua J, Lee HH, Lucas M, Belisle J, Deretic V. Mechanism of phagolysosome biogenesis block by viable Mycobacterium tuberculosis. Proceedings of the National Academy of Sciences of the United States of America. 2005;102(11):4033–8. doi: 10.1073/pnas.0409716102. PubMed PMID: 15753315; PubMed Central PMCID: PMC554822.

40. Hernandez LD, Hueffer K, Wenk MR, Galan JE. Salmonella modulates vesicular traffic by altering phosphoinositide metabolism. Science. 2004;304(5678): 1805–7. doi: 10.1126/science.1098188. PubMed PMID: 15205533.

41. Toulabi L, Wu X, Cheng Y, Mao Y. Identification and structural characterization of a Legionella phosphoinositide phosphatase. The Journal of biological chemistry. 2013;288(34):24518–27. doi: 10.1074/jbc.M113.474239. PubMed PMID: 23843460; PubMed Central PMCID: PMC3750150.

42. Stein MP, Muller MP, Wandinger-Ness A. Bacterial pathogens commandeer Rab GTPases to establish intracellular niches. Traffic. 2012;13(12): 1565–88. doi: 10.1111/tra.12000. PubMed PMID: 22901006; PubMed Central PMCID: PMC3530015.

43. Bore E, Langsrud S, Langsrud O, Rode TM, Holck A. Acid-shock responses in Staphylococcus aureus investigated by global gene expression analysis. Microbiology. 2007;153(Pt 7):2289–303. doi: 10.1099/mic.0.2007/005942-0. PubMed PMID: 17600073.

44. Schwartz JT, Allen LA. Role of urease in megasome formation and Helicobacter pylori survival in macrophages. Journal of leukocyte biology. 2006;79(6): 1214–25. doi: 10.1189/jlb.0106030. PubMed PMID: 16543403; PubMed Central PMCID: PMC1868427.

45. Kohler LJ, Roy CR. Biogenesis of the lysosome-derived vacuole containing Coxiella burnetii. Microbes and infection. 2015;17(11-12):766–71. doi: 10.1016/j.micinf.2015.08.006. PubMed PMID: 26327296; PubMed Central PMCID: PMC4666725.

46. Bera A, Herbert S, Jakob A, Vollmer W, Gotz F. Why are pathogenic staphylococci so lysozyme resistant? The peptidoglycan O-acetyltransferase OatA is the major determinant for lysozyme resistance of Staphylococcus aureus. Molecular microbiology. 2005;55(3):778–87. doi: 10.1111/j.1365-2958.2004.04446.x. PubMed PMID: 15661003.

47. Jin T, Bokarewa M, Foster T, Mitchell J, Higgins J, Tarkowski A. Staphylococcus aureus resists human defensins by production of staphylokinase, a novel bacterial evasion mechanism. Journal of immunology. 2004; 172(2): 1169–76. doi: 10.4049/jimmunol.172.2.1169. PubMed PMID: 14707093.

48. Peschel A, Jack RW, Otto M, Collins LV, Staubitz P, Nicholson G, et al. Staphylococcus aureus resistance to human defensins and evasion of neutrophil killing via the novel virulence factor MprF is based on modification of membrane lipids with l-lysine. The Journal of experimental medicine. 2001;193(9):1067–76. doi: 10.1084/jem.193.9.1067. PubMed PMID: 11342591; PubMed Central PMCID: PMC2193429.

49. Sieprawska-Lupa M, Mydel P, Krawczyk K, Wojcik K, Puklo M, Lupa B, et al. Degradation of human antimicrobial peptide LL-37 by Staphylococcus aureus-derived proteinases. Antimicrobial agents and chemotherapy. 2004;48(12):4673–9. doi: 10.1128/AAC.48.12.4673-4679.2004. PubMed PMID: 15561843; PubMed Central PMCID: PMC529204.

50. Liu GY, Essex A, Buchanan JT, Datta V, Hoffman HM, Bastian JF, et al. Staphylococcus aureus golden pigment impairs neutrophil killing and promotes virulence through its antioxidant activity. The Journal of experimental medicine. 2005;202(2):209–15. doi: 10.1084/jem.20050846. PubMed PMID: 16009720; PubMed Central PMCID: PMC2213009.

51. Ballal A, Manna AC. Regulation of superoxide dismutase (sod) genes by SarA in Staphylococcus aureus. Journal of bacteriology. 2009;191(10):3301–10. doi: 10.1128/JB.01496-08. PubMed PMID: 19286803; PubMed Central PMCID: PMC2687179.

52. Cosgrove K, Coutts G, Jonsson IM, Tarkowski A, Kokai-Kun JF, Mond JJ, et al. Catalase (KatA) and alkyl hydroperoxide reductase (AhpC) have compensatory roles in peroxide stress resistance and are required for survival, persistence, and nasal colonization in Staphylococcus aureus. Journal of bacteriology. 2007;189(3):1025–35. doi: 10.1128/JB.01524-06. PubMed PMID: 17114262; PubMed Central PMCID: PMC1797328.

53. Goncalves VL, Nobre LS, Vicente JB, Teixeira M, Saraiva LM. Flavohemoglobin requires microaerophilic conditions for nitrosative protection of Staphylococcus aureus. FEBS letters. 2006;580(7):1817–21. doi: 10.1016/j.febslet.2006.02.039. PubMed PMID: 16516202.

54. Gomez-Mejia A, Gamez G, Hammerschmidt S. Streptococcus pneumoniae two-component regulatory systems: The interplay of the pneumococcus with its environment. International journal of medical microbiology: IJMM. 2018;308(6):722–37. doi: 10.1016/j.ijmm.2017.11.012. PubMed PMID: 29221986.

55. Mascher T, Heintz M, Zahner D, Merai M, Hakenbeck R. The CiaRH system of Streptococcus pneumoniae prevents lysis during stress induced by treatment with cell wall inhibitors and by mutations in pbp2x involved in beta-lactam resistance. Journal of bacteriology. 2006;188(5):1959–68. doi: 10.1128/JB.188.5.1959-1968.2006. PubMed PMID: 16484208; PubMed Central PMCID: PMC1426552.

56. Ibrahim YM, Kerr AR, McCluskey J, Mitchell TJ. Role of HtrA in the virulence and competence of Streptococcus pneumoniae. Infection and immunity. 2004;72(6):3584–91. doi: 10.1128/IAI.72.6.3584-3591.2004. PubMed PMID: 15155668; PubMed Central PMCID: PMC415679.

57. Hentrich K, Lofling J, Pathak A, Nizet V, Varki A, Henriques-Normark B. Streptococcus pneumoniae Senses a Human-like Sialic Acid Profile via the Response Regulator CiaR. Cell host & microbe. 2016;20(3):307–17. doi: 10.1016/j.chom.2016.07.019. PubMed PMID: 27593514; PubMed Central PMCID: PMC5025396.

58. Ibrahim YM, Kerr AR, McCluskey J, Mitchell TJ. Control of virulence by the two-component system CiaR/H is mediated via HtrA, a major virulence factor of Streptococcus pneumoniae. Journal of bacteriology. 2004;186(16):5258–66. doi: 10.1128/JB.186.16.5258-5266.2004. PubMed PMID: 15292127; PubMed Central PMCID: PMC490881.

59. Spiess C, Beil A, Ehrmann M. A temperature-dependent switch from chaperone to protease in a widely conserved heat shock protein. Cell. 1999;97(3):339–47. doi: 10.1016/s0092-8674(00)80743-6. PubMed PMID: 10319814.

60. Marx P, Meiers M, Bruckner R. Activity of the response regulator CiaR in mutants of Streptococcus pneumoniae R6 altered in acetyl phosphate production. Frontiers in microbiology. 2014;5:772. doi: 10.3389/fmicb.2014.00772. PubMed PMID: 25642214; PubMed Central PMCID: PMC4295557.

61. Mahdi LK, Wang H, Van der Hoek MB, Paton JC, Ogunniyi AD. Identification of a novel pneumococcal vaccine antigen preferentially expressed during meningitis in mice. The Journal of clinical investigation. 2012;122(6):2208–20. doi: 10.1172/JCI45850. PubMed PMID: 22622042; PubMed Central PMCID: PMC3366392.

62. Lam GY, Huang J, Brumell JH. The many roles of NOX2 NADPH oxidase-derived ROS in immunity. Seminars in immunopathology. 2010;32(4):415–30. doi: 10.1007/s00281-010-0221-0. PubMed PMID: 20803017.

63. Turk V, Stoka V, Vasiljeva O, Renko M, Sun T, Turk B, et al. Cysteine cathepsins: from structure, function and regulation to new frontiers. Biochimica et biophysica acta. 2012;1824(1):68–88. doi: 10.1016/j.bbapap.2011.10.002. PubMed PMID: 22024571; PubMed Central PMCID: PMC7105208.

64. Echlin H, Frank MW, Iverson A, Chang TC, Johnson MD, Rock CO, et al. Pyruvate Oxidase as a Critical Link between Metabolism and Capsule Biosynthesis in Streptococcus pneumoniae. PLoS pathogens. 2016;12(10):e1005951. doi: 10.1371/journal.ppat.1005951. PubMed PMID: 27760231; PubMed Central PMCID: PMC5070856.

65. Pericone CD, Park S, Imlay JA, Weiser JN. Factors contributing to hydrogen peroxide resistance in Streptococcus pneumoniae include pyruvate oxidase (SpxB) and avoidance of the toxic effects of the fenton reaction. Journal of bacteriology. 2003;185(23):6815–25. doi: 10.1128/jb.185.23.6815-6825.2003. PubMed PMID: 14617646; PubMed Central PMCID: PMC262707.

66. Pericone CD, Overweg K, Hermans PW, Weiser JN. Inhibitory and bactericidal effects of hydrogen peroxide production by Streptococcus pneumoniae on other inhabitants of the upper respiratory tract. Infection and immunity. 2000;68(7):3990–7. doi: 10.1128/iai.68.7.3990-3997.2000. PubMed PMID: 10858213; PubMed Central PMCID: PMC101678.

67. Rai P, Parrish M, Tay IJ, Li N, Ackerman S, He F, et al. Streptococcus pneumoniae secretes hydrogen peroxide leading to DNA damage and apoptosis in lung cells. Proceedings of the National Academy of Sciences of the United States of America. 2015;112(26):E3421–30. doi: 10.1073/pnas.1424144112. PubMed PMID: 26080406; PubMed Central PMCID: PMC4491788.

68. Braun JS, Sublett JE, Freyer D, Mitchell TJ, Cleveland JL, Tuomanen EI, et al. Pneumococcal pneumolysin and H(2)O(2) mediate brain cell apoptosis during meningitis. The Journal of clinical investigation. 2002;109(1):19–27. doi: 10.1172/JCI12035. PubMed PMID: 11781347; PubMed Central PMCID: PMC150815.

69. Hoffmann O, Zweigner J, Smith SH, Freyer D, Mahrhofer C, Dagand E, et al. Interplay of pneumococcal hydrogen peroxide and host-derived nitric oxide. Infection and immunity. 2006;74(9):5058–66. doi: 10.1128/IAI.01932-05. PubMed PMID: 16926397; PubMed Central PMCID: PMC1594840.

70. Joshi N, Caputo GM, Weitekamp MR, Karchmer AW. Infections in patients with diabetes mellitus. The New England journal of medicine. 1999;341(25): 1906–12. doi: 10.1056/NEJM199912163412507. PubMed PMID: 10601511.

71. van Veen KE, Brouwer MC, van der Ende A, van de Beek D. Bacterial meningitis in diabetes patients: a population-based prospective study. Scientific reports. 2016;6:36996. doi: 10.1038/srep36996. PubMed PMID: 27845429; PubMed Central PMCID: PMC5109544.

72. Pomar V, de Benito N, Mauri A, Coll P, Gurgui M, Domingo P. Characteristics and outcome of spontaneous bacterial meningitis in patients with diabetes mellitus. BMC infectious diseases. 2020;20(1):292. doi: 10.1186/s12879-020-05023-5. PubMed PMID: 32312231; PubMed Central PMCID: PMC7171854.

73. Thomsen RW, Hundborg HH, Lervang HH, Johnsen SP, Sorensen HT, Schonheyder HC. Diabetes and outcome of community-acquired pneumococcal bacteremia: a 10-year population-based cohort study. Diabetes care. 2004;27(1):70–6. doi: 10.2337/diacare.27.1.70. PubMed PMID: 14693969.

74. Stins MF, Badger J, Sik Kim K. Bacterial invasion and transcytosis in transfected human brain microvascular endothelial cells. Microbial pathogenesis. 2001;30(1):19–28. doi: 10.1006/mpat.2000.0406. PubMed PMID: 11162182.

75. Hammerschmidt S, Wolff S, Hocke A, Rosseau S, Muller E, Rohde M. Illustration of pneumococcal polysaccharide capsule during adherence and invasion of epithelial cells. Infection and immunity. 2005;73(8):4653–67. doi: 10.1128/IAI.73.8.4653-4667.2005. PubMed PMID: 16040978; PubMed Central PMCID: PMC1201225.

76. Dixit C, Keller LE, Bradshaw JL, Robinson DA, Swiatlo E, McDaniel LS. Nonencapsulated Streptococcus pneumoniae as a cause of chronic adenoiditis. IDCases. 2016;4:56–8. doi: 10.1016/j.idcr.2016.04.001. PubMed PMID: 27144125; PubMed Central PMCID: PMC4840421.

77. Bradshaw JL, Pipkins HR, Keller LE, Pendarvis JK, McDaniel LS. Mucosal Infections and Invasive Potential of Nonencapsulated Streptococcus pneumoniae Are Enhanced by Oligopeptide Binding Proteins AliC and AliD. mBio. 2018;9(1). doi: 10.1128/mBio.02097-17. PubMed PMID: 29339428; PubMed Central PMCID: PMC5770551.

78. Keller LE, Robinson DA, McDaniel LS. Nonencapsulated Streptococcus pneumoniae: Emergence and Pathogenesis. mBio. 2016;7(2):e01792. doi: 10.1128/mBio.01792-15. PubMed PMID: 27006456; PubMed Central PMCID: PMC4807366.

79. Bradshaw JL, Rafiqullah IM, Robinson DA, McDaniel LS. Transformation of nonencapsulated Streptococcus pneumoniae during systemic infection. Scientific reports. 2020;10(1):18932. doi: 10.1038/s41598-020-75988-5. PubMed PMID: 33144660; PubMed Central PMCID: PMC7641166.

80. Badgujar DC, Anil A, Green AE, Surve MV, Madhavan S, Beckett A, et al. Structural insights into loss of function of a pore forming toxin and its role in pneumococcal adaptation to an intracellular lifestyle. PLoS pathogens. 2020;16(11):e1009016. doi: 10.1371/journal.ppat.1009016. PubMed PMID: 33216805.

81. Sorg RA, Kuipers OP, Veening JW. Gene expression platform for synthetic biology in the human pathogen Streptococcus pneumoniae. ACS synthetic biology. 2015;4(3):228–39. doi: 10.1021/sb500229s. PubMed PMID: 24845455.

82. Kjos M, Aprianto R, Fernandes VE, Andrew PW, van Strijp JA, Nijland R, et al. Bright fluorescent Streptococcus pneumoniae for live-cell imaging of host-pathogen interactions. Journal of bacteriology. 2015;197(5):807–18. doi: 10.1128/JB.02221-14. PubMed PMID: 25512311; PubMed Central PMCID: PMC4325099.

