## Supplementary figures and images for "Pyruvate oxidase as a key determinant of pneumococcal viability during transcytosis across the blood-brain barrier endothelium"

### Contribution of hydrogen peroxide to pneumococcal survival within the vacuoles of brain endothelial cells

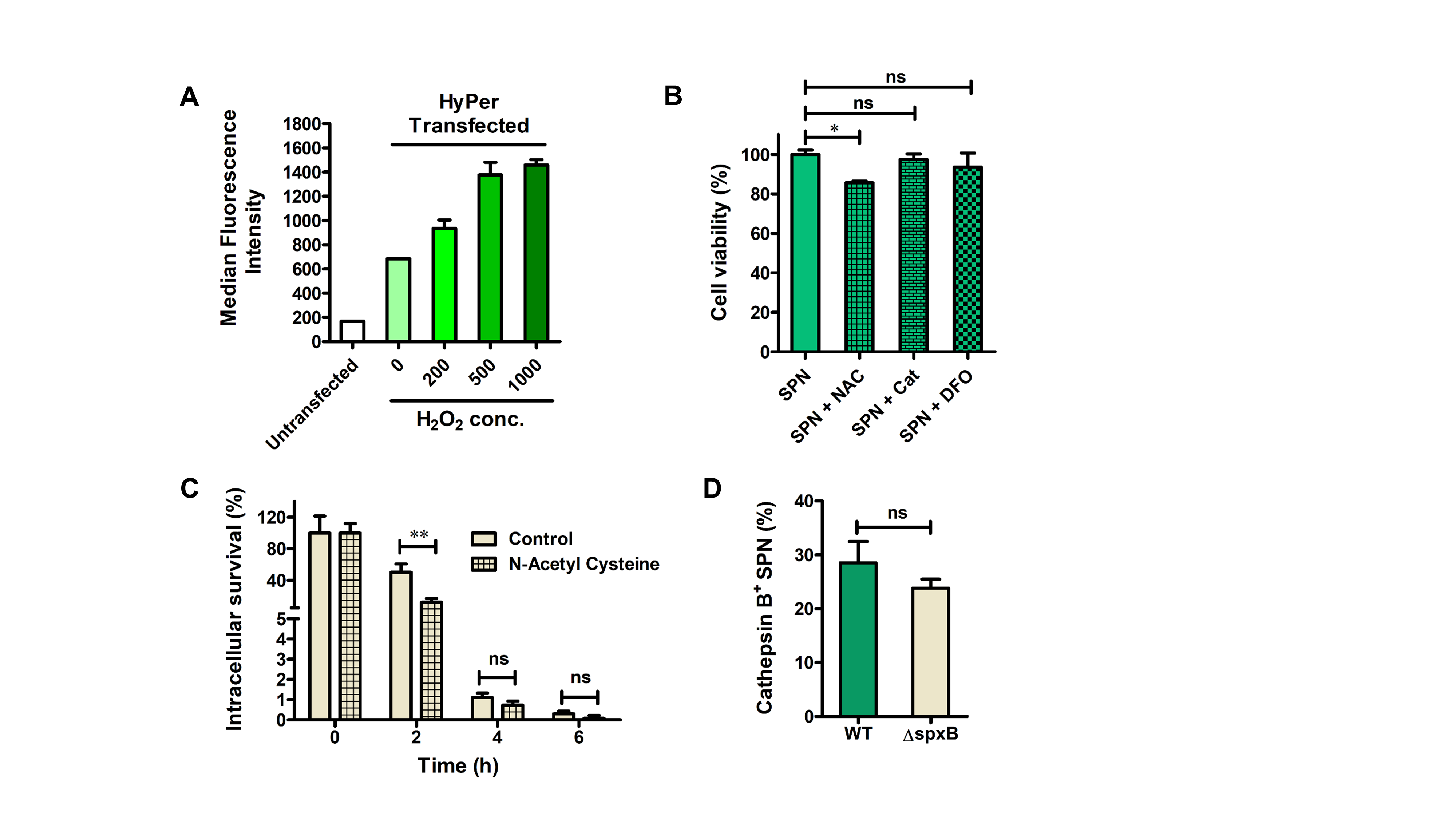

### Pyruvate oxidase modulates pneumococcal interaction with the brain endothelial cells

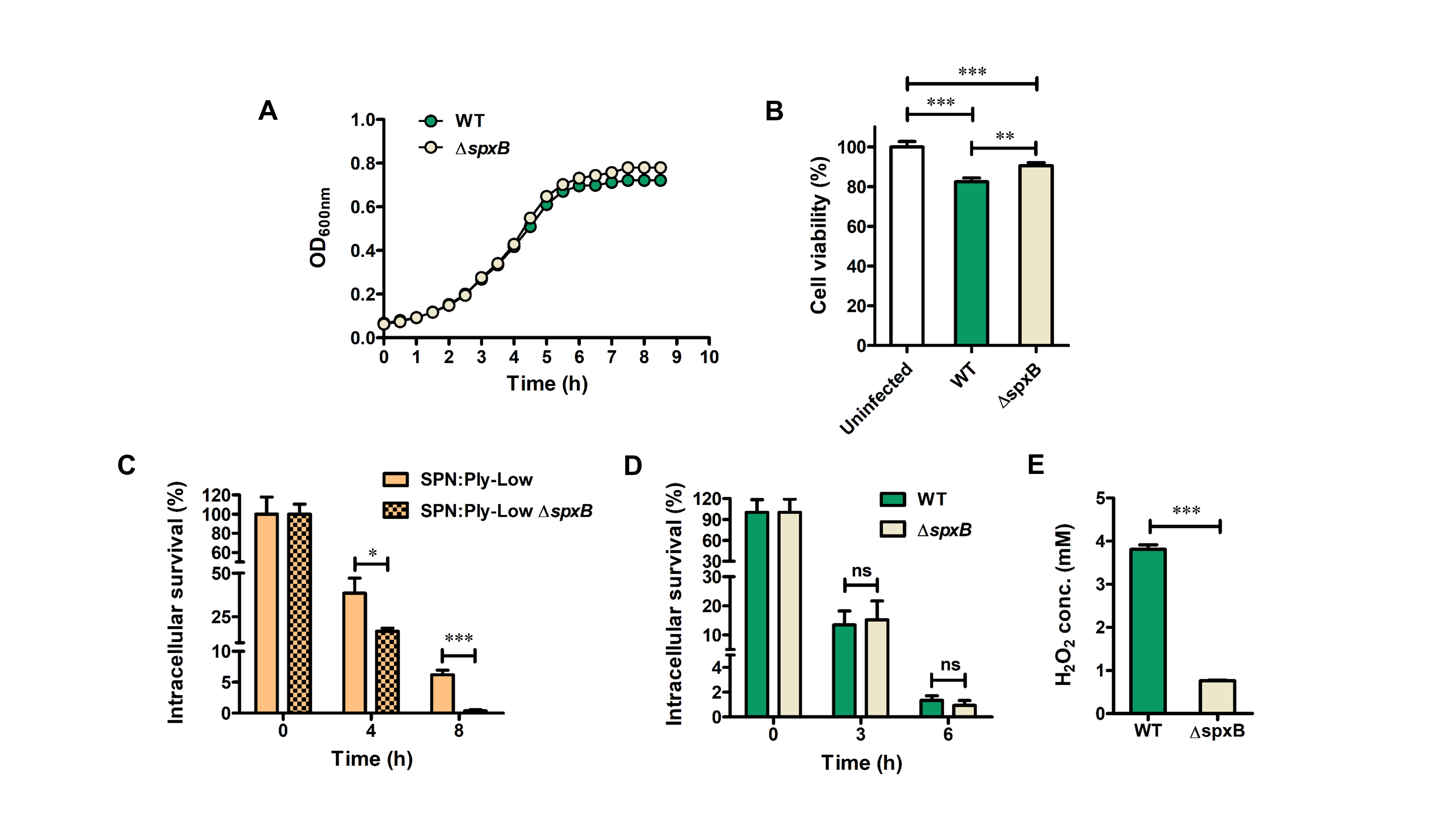

### Relative contribution of CiaRH and pyruvate oxidase towards pneumococcal tolerance to acidic stress

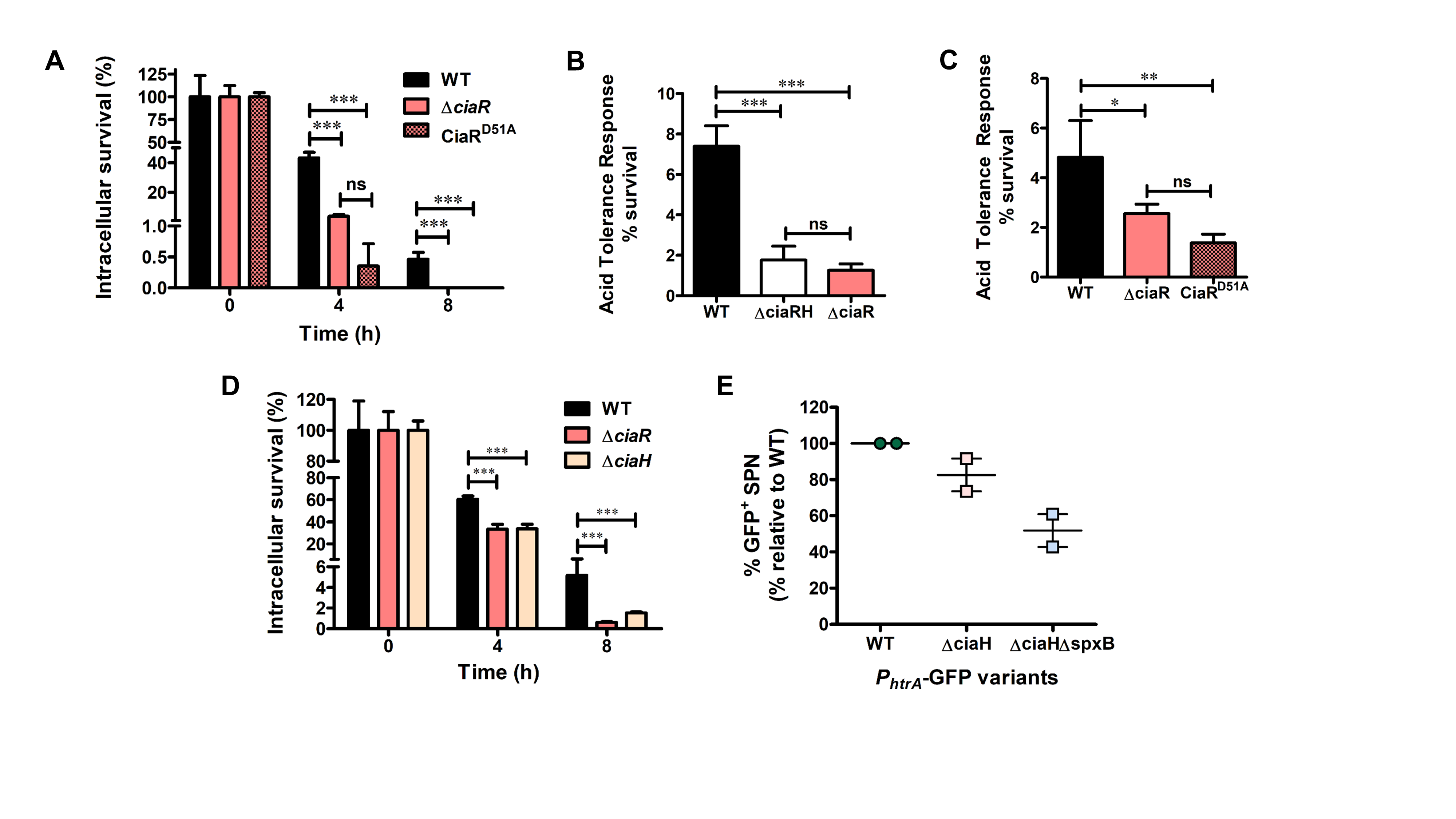
