## Supplementary material for "Pyruvate oxidase as a key determinant of pneumococcal viability during transcytosis across the blood-brain barrier endothelium": List of strains

**S1 Table**

| **Strains** | **Nomenclature** | **Source** |
| --- | --- | --- |
| R6 |  | Prof. TJ Mitchell, Univ. of Birmingham, UK |
| R6Δ*spxB* | ABP101 | This study |
| R6:*hlpA*-GFP/tagRFP | ABP107/ABP109 | This study |
| R6Δ*spxB*:*hlpA*-GFP/tagRFP | ABP111/ABP110 | This study |
| R6Δ*ply* | ABP007 | Badgujar *et al* [80] |
| R6Δ*ply*Δ*spxB* | ABP104 | This study |
| R6:Ply-Low | ABP015 | Surve *et al* [13] |
| R6:Ply-LowΔ*spxB* | ABP130 | This study |
| R6Δ*ciaR* | ABP121 | This study |
| R6Δ*ciaH* | ABP122 | This study |
| R6Δ*ciaRH* | ABP128 | This study |
| R6:CiaR^D51A^ | ABP133 | This study |
| R6Δ*ciaH*Δ*spxB* | ABP124 | This study |
| R6:*P_htrA_*-GFP | ABP136 | This study |
| R6:*P_htrA_*-GFPΔ*ciaH* | ABP137 | This study |
| R6:*P_htrA_*-GFPΔ*ciaH*Δ*spxB* | ABP138 | This study |
| D39 |  | Prof EI Tuomanen, St. Jude Childrens Hospital, USA |
| D39Δ*spxB* | ABP135 | This study |
| TIGR4 |  | Prof TJ Mitchell, Univ. of Birmingham, UK |
| TIGR4Δ*spxB* | ABP102 | This study |
