## Supplementary material for "Pyruvate oxidase as a key determinant of pneumococcal viability during transcytosis across the blood-brain barrier endothelium": List of primers

**S2 Table**

| **Sl No.** | **Name** | **5’ to 3’ sequence** |
| --- | --- | --- |
| 1 | spxB-F1 | CTGTTCTGACCAGTTCCCTT |
| 2 | spxB-R1 | TCCAAGATATGCTCCAAGTC |
| 3 | spxB-F2-R6 | GACGGTATCAATGACGCTCC |
| 4 | spxB-R2 | TCAAGCACTAGACCAGCAGG |
| 5 | ciaRdel-up-F | ataatgCTCGAGCTTGATACTTATACTCACGC |
| 6 | ciaRdel-up-R | aattctaGAATTCTTATGCATTTCCGTATTGAAGA |
| 7 | ciaR-5UTR-F | aattctaGGATCCATAAGCCTAAAATAAAAAGAAAACTC |
| 8 | ciaR-5UTR-R | aagtttactgaaCATGAGAAACTCCTCCTTATTAAAA |
| 9 | ciaRdel-down-F | gaggagtttctcATGTTCAGTAAACTTAAAAAAACAT |
| 10 | ciaRdel-down-R | atgctatTCTAGAATCCAAAATATCGCCCCAATTG |
| 11 | ciaHdel-up-F | ataatgCTCGAGATAAGCCTAAAATAAAAAGAAAACTC |
| 12 | ciaHdel-up-R | aattctaGAATTCAAAGTCATCCGCATACCATGT |
| 13 | ciaHdel-down-F | aattctaGGATCCAAATATCGCTCCAATTGGGGCG |
| 14 | ciaHdel-down-R | atgctatTCTAGAACTTTCAGCAGTTGCTGTAATGAC |
| 15 | ciaRH-F-EcoRI | aattctaGAATTCATAAGCCTAAAATAAAAAGAAAACTC |
| 16 | ciaRH-R-BamHI | aattctaGGATCCATCCAAAATATCGCCCCAATTG |
| 17 | SDM-D51A-F | TGATTTTGCTGGCTTTGATGTTGCCAGAAAAAAATGGTT |
| 18 | SDM-D51A-R | GGCAACATCAAAGCCAGCAAAATCAAGTCATAGACACCA |
| 19 | ciaRH-Flnk-F | CGGAGAATCAGATGAGGATGAA |
| 20 | ciaRH-Flnk-R | GTCTAAGAAAGGCTTGATACGG |
| 21 | PhtrA-F2-XbaI | ATGCTATTCTAGACACATCTTATTCACAAAATA |
| 22 | PhtrA-R2-BamHI | AATTCTAGGATCCCATATTTGCCTCCATATGTT |
| 23 | pPEP-Flnk-F | CCAACCTAACCAGCTACCAAG |
| 24 | pPEP-Flnk-R | CATGGCACGGCTAAGATGTTG |
| 25 | lox-HpaI-Fo | agccttGTTAACTACCGTTCGTATAGCATACATTA |
| 26 | lox-BamHI-Rn | gtaataGGATCCGGAGCTCTCCCATATGGTCG |
| 27 | Cm-pPEP-EcoRI-F | atgctatGAATTCGCACCCATTAGTTCAACAAA |
| 28 | Cm-BamHI-R | atccGGATCCTACAGTCGGCATTATCTCATA |
| 29 | spec-EcoRI-F | atgctatGAATTCCATATATAATCTAGAATAAAATTAAC |
| 30 | spec-R-BamHI | atccGGATCCAATCTGATTACCAATTAGAATG |
| 31 | hlpA-up-F | AACAAGTCAGCCACCTGTAG |
| 32 | hlpA-down-R | CGTGGCTGACGATAATGAGG |
| 33 | hlpA-flanking-F | AAAGGTGATGAGGGTAATGCCTGTC |
| 34 | hlpA-flanking-R | GTAACCAACCTCAACATTAGCGCC |

*Restriction sites are shown in red.
